# The structure of the apo-PIWI HSP90 complex

**DOI:** 10.64898/2026.07.04.736496

**Authors:** Peter Donlon, Paula Sotelo-Parrilla, David MacKenzie MacLeod, Aleksandra Rosinska, Tamoghna Chowdhury, Katherine Isabella Leith, Ansgar Zoch, Christos Spanos, Atlanta G. Cook, A. Arockia Jeyaprakash, Dónal O’Carroll

## Abstract

PIWI proteins are members of the Argonaute family and together with piRNAs protect metazoan germlines from transposons. PIWI proteins adopt a bi-lobed architecture with a central RNA-binding channel. HSP90 function has been linked to piRNA biogenesis, but the precise molecular mechanism is unresolved. Using the mammalian embryonic piRNA pathway as a model system, we find compelling evidence for the existence of PIWIL2- (MILI-) and PIWIL4- (MIWI2-) HSP90 complexes in foetal testis. We purify apo-PIWIL4-HSP90 from cells and determine its structure by cryo-electron microscopy. Distinct from piRNA-bound PIWI, apo-PIWIL4 adopts a unique and open conformation. The HSP90 dimer binds and unfolds PIWI’s linker 1 domain. PIWIL4’s N domain and the RNA-binding PAZ-MID-PIWI module are placed on opposite sides of the HSP90 dimer’s lumen. We further demonstrate that PIWI-HSP90 complexes, the open apo-PIWI conformation, and the HSP90 lumen-binding peptide are conserved features of PIWI proteins.

## Introduction

piRNA biogenesis and PIWI loading are central to anti-transposon immunity that protects the continuity of the germline. In mammals, genome methylation is erased and reset during germ cell development. The loss of DNA methylation unleashes transposon expression which poses an existential threat to the germline. The PIWI protein PIWIL2 (MILI) is a cytoplasmic piRNA-guided slicer endonuclease that cleaves transposon transcripts, preventing transposition^1,2^. piRNAs also identify active transposon loci by tethering the PIWI protein PIWIL4 (MIWI2) to their nascent transcripts which leads to the restoration of epigenetic transposon silencing through piRNA-directed DNA methylation^1–3^. Several types of piRNA biogenesis pathways exist to produce, amplify and diversify the piRNA repertoire^2,4–10^. Each of these pathways face a common challenge: the single-stranded precursor-piRNA (pre-piRNA) needs to be incorporated into PIWI proteins that have closed structures.

PIWIs have a typical Argonaute protein domain composition consisting of N, PAZ, MID and PIWI domains that fold to form a bi-lobed architecture with a central RNA-binding channel^11–18^. The 5’ phosphate of the piRNA is bound by the MID domain in the MID-PIWI lobe and the 3’ end is bound by the PAZ domain in the N-PAZ lobe^11–21^. The piRNA is threaded through the central channel and interacts with all domains. How the single-stranded pre-piRNA is incorporated into this complex bi-lobed structure remains unknown. HSP70/90 chaperone function has been linked to Argonaute-mediated small RNA pathways in animals, plants and yeast^22–38^. HSP70/90 chaperones have known roles in mediating ligand binding through conformational change of their clients^39^. For example, the glucocorticoid receptor requires HSP-mediated conformational change to bind glucocorticoid^40^. In this process, HSP70 with the co-chaperone STIP1 loads two HSP90s onto the glucocorticoid receptor. ATP binding to HSP90, displacement of HSP70, replacement of STIP1 with co-chaperones P23 and FKBP4 result in a glucocorticoid receptor conformation that is competent to bind glucocorticoid with high affinity^41–43^. Very recent structural studies revealed the AGO2 maturation complex^44^. In this structure, HSP90 and the co-chaperone P23 open AGO2, where N domain and the RNA binding module (PAZ–MID–PIWI domains) are fully detached and placed on opposite sides of the HSP90 dimer^44^. This conformation exposes a positively charged cleft that accommodates a duplex RNA within the 5’ and 3’ RNA-binding pockets within AGO2^44^. In our study, we sought to understand the mechanism of PIWI protein loading with pre-piRNAs.

## Results

### PIWIL2 and PIWIL4 associate with chaperones in foetal germ cells

To identify factors that are required for reconfiguring apo-PIWI proteins, we chose to study the embryonic piRNA pathway at embryonic day 16.5 (E16.5) for several reasons. Firstly, primary and secondary piRNA biogenesis pathways are active^1,2,10^. Secondly, PIWIL2 receives piRNAs from all biogenesis pathways whereas PIWIL4 receives mostly secondary piRNAs^10,45^. Thirdly, it is a timepoint when a substantial fraction of PIWIL4 is cytoplasmic and engaged in piRNA biogenesis^2,46^. While PIWIL2 and PIWIL4 may associate with different biogenesis factors and effectors, the factors required for reconfiguring apo-PIWI proteins to load pre-piRNA should be common to both. Using the *Piwil4^HA^* mouse allele^45^, we had previously characterised PIWIL4-associated proteins in foetal testes^46^ (Figure 1A). To compare PIWIL2 and PIWIL4-associated proteins in E16.5 foetal gonocytes, we generated a *Piwil2^HA^* mouse allele, which encodes a fully functional N-terminal HA epitope PIWIL2 protein (Figure S1A-G) and performed immunoprecipitation coupled with quantitative mass spectrometry (IP-MS) of HA-PIWIL2 (Figure 1B). In the overlap between PIWIL2 and PIWIL4-associated proteins from foetal testes, many piRNA biogenesis factors were present as expected, but also notable was the presence of many chaperones and co-chaperones (Figure 1C). HSP90α and HSP90β associated with PIWIL2 and PIWIL4, as did HSC70 (the constitutively expressed variant of HSP70) and the co-chaperones STIP1, P23 (PTGES3), FKBP4 and FKBP6. FKBP6 is essential for secondary piRNA biogenesis^29^ and HSP90α for mouse spermatogenesis^30^. To validate our data, we firstly performed IP-MS of HSP90α from E16.5 foetal testes. HSP90α associated with PIWIL2 and PIWIL4 as well as many piRNA biogenesis factors (Figure 1D). We next performed co-IF of HA-PIWIL4 with HSP90α, HSP90β and HSC70 in *Piwil4^HA/+^* foetal gonocytes. piRNA biogenesis occurs in the cytoplasm in nuage^47^. In the cytoplasm HA-PIWIL4 colocalised to a large degree with HSP90α, HSP90β and HSC70 (Figure 1E and Figure S1H). HA-PIWIL2 also partially co-localised with HSP90α, HSP90β and HSC70 in *Piwil2^HA/+^* foetal gonocytes (Figure 1F and Figure S1I). Given our data and the fact that chaperone function has also been linked to Argonaute-mediated small RNA pathways in animals, plants and yeast^22–38^, we selected chaperones for further analyses.

**Figure 1:**
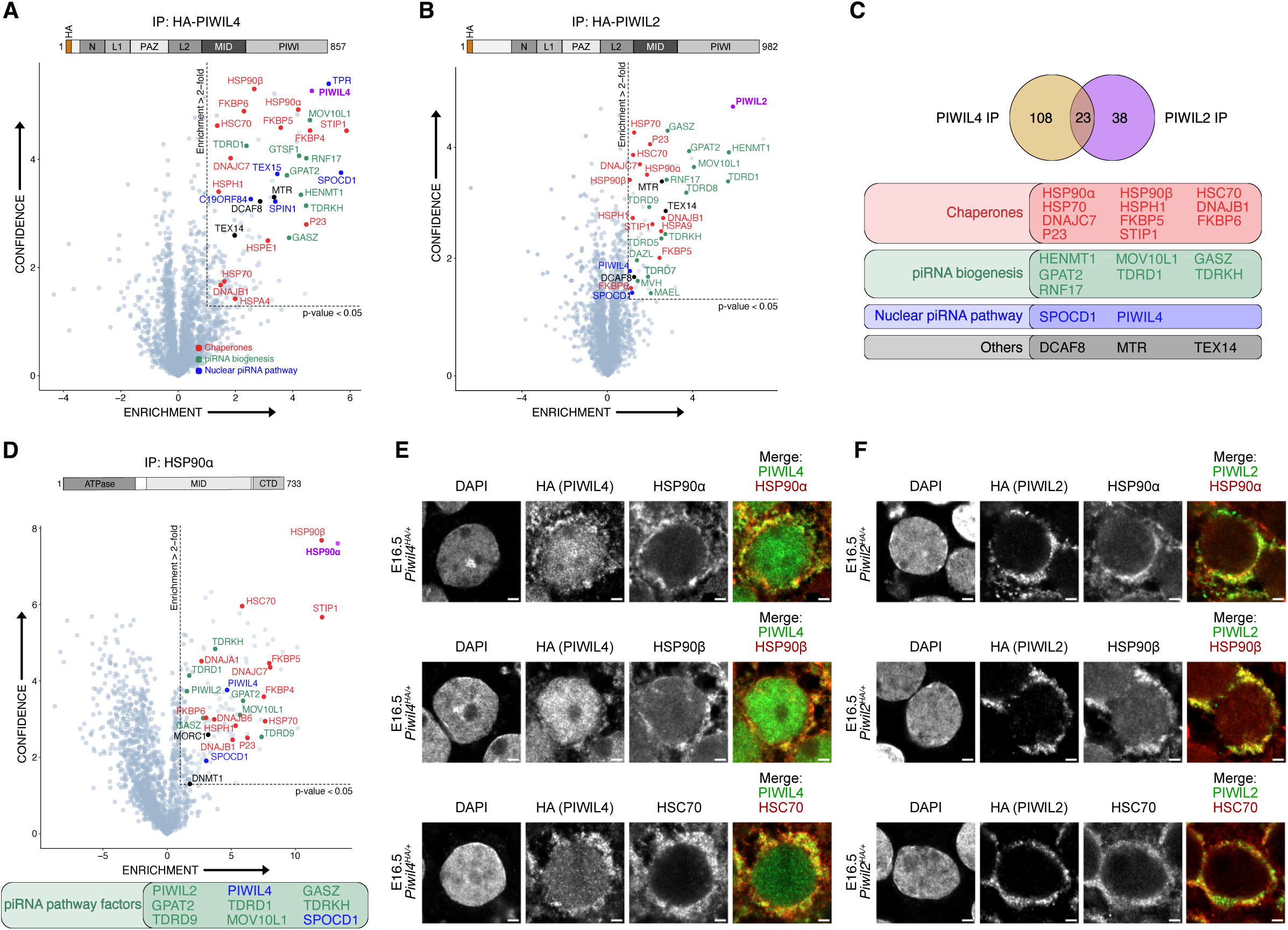
Mouse PIWIL4 and PIWIL2 associate with the HSP90/70 chaperone machinery in foetal testes **A-B**, Volcano plot showing enrichment (log_2_(mean LFQ ratio of anti-HA immunoprecipitates from *Piwil4^HA/HA^* (**A**) or *Piwil2^HA/+^* (**B**) vs wild-type E16.5 foetal testis lysates) and statistical confidence of proteins co-purifying with HA-PIWIL4 (**A**, n = 3 with 50 testes per replicate, reanalysed from previously published data^46^) or HA-PIWIL2 (**B**, n = 3 with 50 testes per replicate). **C**, Schematic representation of the overlapping set of proteins co-purifying with both PIWIL4 and PIWIL2. In **A-C**, the immunoprecipitation target is indicated in purple, piRNA biogenesis related factors in green, chaperones and co-chaperones in red, and nuclear piRNA-directed DNA methylation factors in blue. Other proteins are not highlighted. **D**, Volcano plot showing enrichment (log_2_(mean LFQ ratio of anti-HSP90α immunoprecipitates/anti-rabbit serum immunoprecipitated from wild-type E16.5 foetal testis lysates) and statistical confidence of proteins co-purifying with HSP90α (n = 3 with 25 foetal testes per replicate). Known piRNA-related proteins co-purifying with HSP90α are listed below the plot. **E-F**, Representative E16.5 gonocytes stained for HA (HA-PIWIL4 in (**E**) and HA-PIWIL2 in (**F**)), HSP90α (top row), HSP90β (middle row), HSC70 (bottom row) and DAPI in foetal testis sections from *Piwil4^HA/+^* (**E**) or *Piwil2^HA/+^* (**F**) mice. Images are representative of n = 3 biological replicates. Scale bars, 2 μm.

### Purification of apo-PIWIL4-chaperone complexes

If chaperones alter PIWI structure for loading, then it is likely to be a very transient event. To capture this complex, we expressed and purified mouse FLAG-PIWIL4 from mouse embryonic stem cells (ESCs) that lack piRNA biogenesis but express high levels of the HSC70/90 chaperone system. FLAG-PIWIL4 is cytoplasmic in ESCs (Figure 2A). This cytoplasmic localization indicates that FLAG-PIWIL4 is unloaded as it only translocates to the nucleus when it is bound to a piRNA^2^. FLAG-PIWIL4 co-purified with six prominent proteins when resolved on an acrylamide gel and stained with Coomassie (Figure 2B). Mass spectrometry identified these proteins as HSP90α, HSP90β, HSC70 and the co-chaperones STIP1, FKBP4 and P23. The addition of ATP and molybdate to the HSP90-loading complex promotes the formation of HSP90-P23-client and HSP90-FKBP4-client maturation complexes^42,43^. Treatment of the FLAG-PIWIL4 purifications with ATP/molybdate catalyses the same reaction with a dramatic reduction of HSC70 and loss of STIP1 (Figure 2B). In summary, in the absence of precursor-piRNA PIWIL4 co-purifies with chaperones that have established roles in mediating ligand binding through conformational change.

**Figure 2:**
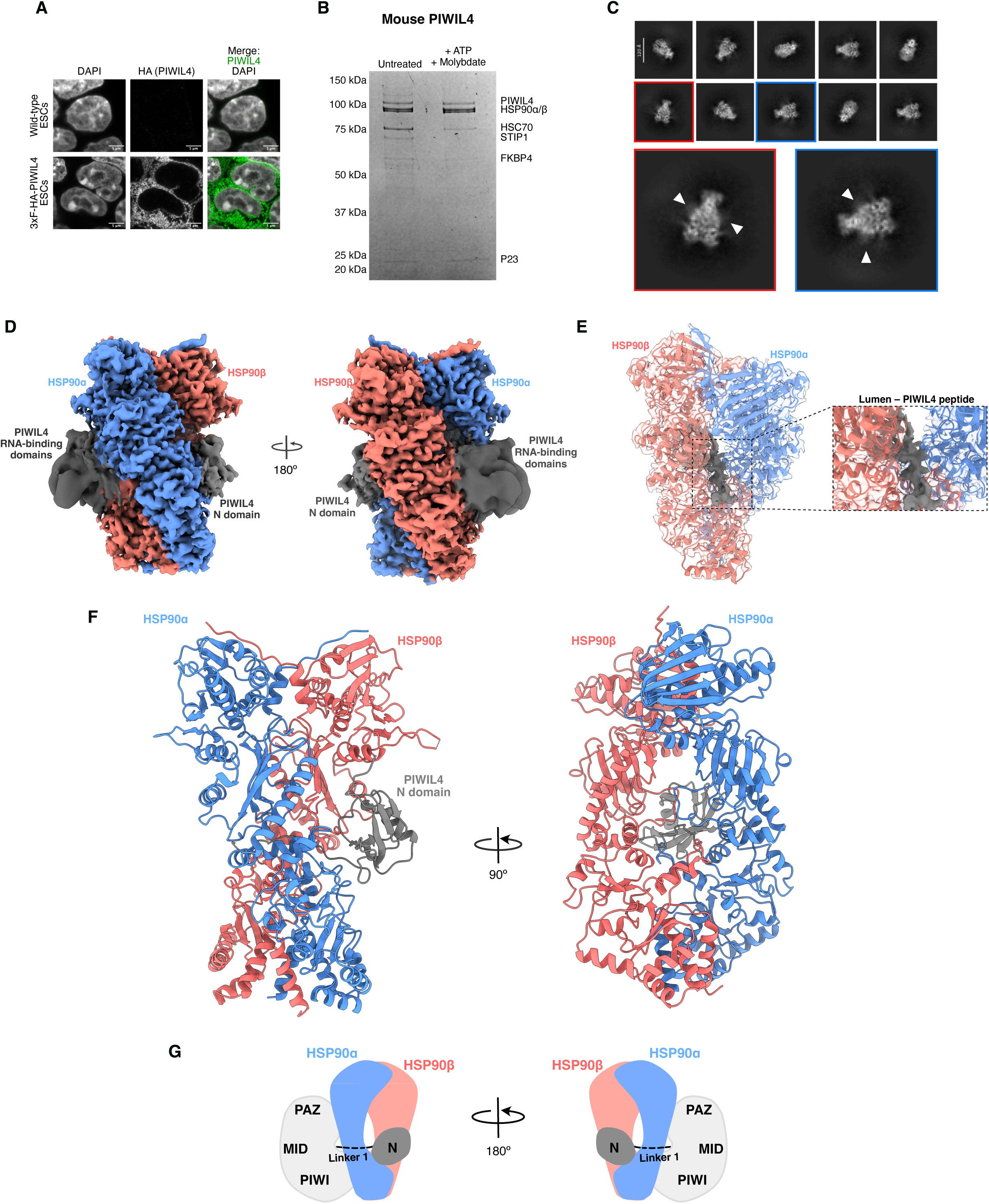
The structure of apo-PIWIL4-HSP90α/β. **A**, Representative images of wild-type, untransfected ESCs (top row) or ESCs stably expressing 3xFLAG-3C-HA-mPIWIL4 (bottom row) stained for HA (HA-PIWIL4) and DAPI. Images are representative of n = 3 independent experiments. Scale bars, 5 μm. **B**, Representative Coomassie-stained SDS-PAGE gel of anti-FLAG immunoprecipitates from 3xFLAG-3C-HA-mPIWIL4-expressing ESCs lysed either in the absence (untreated) or presence of ATP and sodium molybdate. Proteins indicated were identified by band excision and mass spectrometry. For uncropped whole gel source images, see Supplementary Fig. 1Y. **C,** Representative 2D classes of PIWIL4-HSP90 complex, with 2D classes showcasing densities on either side of the lumen as enlarged images and diffuse densities indicated by white arrows **D**, Cryo-EM density map of mouse PIWIL4 (grey) in complex with HSP90α (blue) and HSP90β (salmon) in the HSP90 closed state. **E,** Close-up view of PIWIL4 peptide within the HSP90 dimer lumen. **F,** Cartoon representation of the resolved regions of the mouse PIWIL4-HSP90 structure in two orientations, with the same colours as in **D**. **G**, Schematic of the apo-PIWIL4-HSP90 complex with the well resolved N domain (dark grey) on one side of the HSP90 lumen, with the poorly resolved PAZ-MID-PIWI domains (light grey) on the other side.

### HSP90s reconfigure PIWIL4 into an open conformation

To understand how chaperone engagement remodels PIWIL4 into a conformation suitable to accept precursor-piRNA, we determined the cryo-EM structure of the ATP/molybdate-treated sample at 3.2 Å resolution (Figure 2C-F and Figure S2). Our structure shows a single molecule of mouse PIWIL4 bound to an HSP90 heterodimer composed of HSP90α and HSP90β in the closed conformation (Figure 2D-F). In the closed configuration, the HSP90 middle domains pack closely against one another to form a central lumen that is responsible for client engagement, and the N-terminal and C-terminal domains establish the dimerization interface. We observed a well-defined peptide-like density in the lumen (Figure 2E) with heterogeneous density at either end, which was also visible in 2D class averages (Figure 2C-D). Further focused classification and refinement, using masks covering the heterogeneous densities, improved the density on one side, with dimensions matching those of the PIWIL4 N domain (Figure 2F and Figure S2F). In addition, local filtering of the map revealed an even larger diffused density on the opposite side of the lumen corresponding to the RNA-binding PAZ/MID/PIWI domains. Thus, our structure reveals how HSP90 clamps onto PIWIL4, resulting in an open conformation (Figure 2G).

### The interaction between PIWIL4 and the HSP90 lumen is stabilised by a highly conserved motif

We then sought to define the structural basis of the interaction between PIWIL4 and the HSP90 lumen. The local resolution of the lumen peptide (2.5 - 3.0 Å; Figure S2G) allowed us to unambiguously assign the lumen density to PIWIL4 amino acid residues 203-211, located in Linker 1 (L1) that connects the N domain and the RNA-binding PAZ/MID/PIWI domains (Figure 3A-F). Two basic residues in PIWIL4, Arg206 and Lys207, form salt bridges with HSP90α Glu452 and HSP90β Glu443, respectively (Figure 3B). The flanking PIWIL4 hydrophobic residues Leu203, Ile204, Phe205, Ile208 and Leu209 are buried within two hydrophobic luminal clefts formed by HSP90α Phe353, Leu448, Ile526, Tyr529 and HSP90β Ala610 and Leu611 and HSP90α Ala619, Leu620 and HSP90β Phe344, Leu449, Tyr520 (Figure 3C-F). At the luminal exit, PIWIL4 Lys210 is in close proximity to HSP90α Glu354, suggesting additional electrostatic interactions (Figure 3B). Upon binding to HSP90, the lumen peptide has a buried surface area (BSA) of 2318.3Å^2^, of which 1587.1 Å^2^ correspond to non-polar contacts. This suggests that extensive hydrophobic interactions, together with electrostatic interactions, constitute the primary driving force for HSP90 to stabilise PIWIL4 in an open conformation, consistent with the established role of HSP90 in clamping onto a combined hydrophobic and polar region of client proteins during maturation^41–44,48,49^.

**Figure 3:**
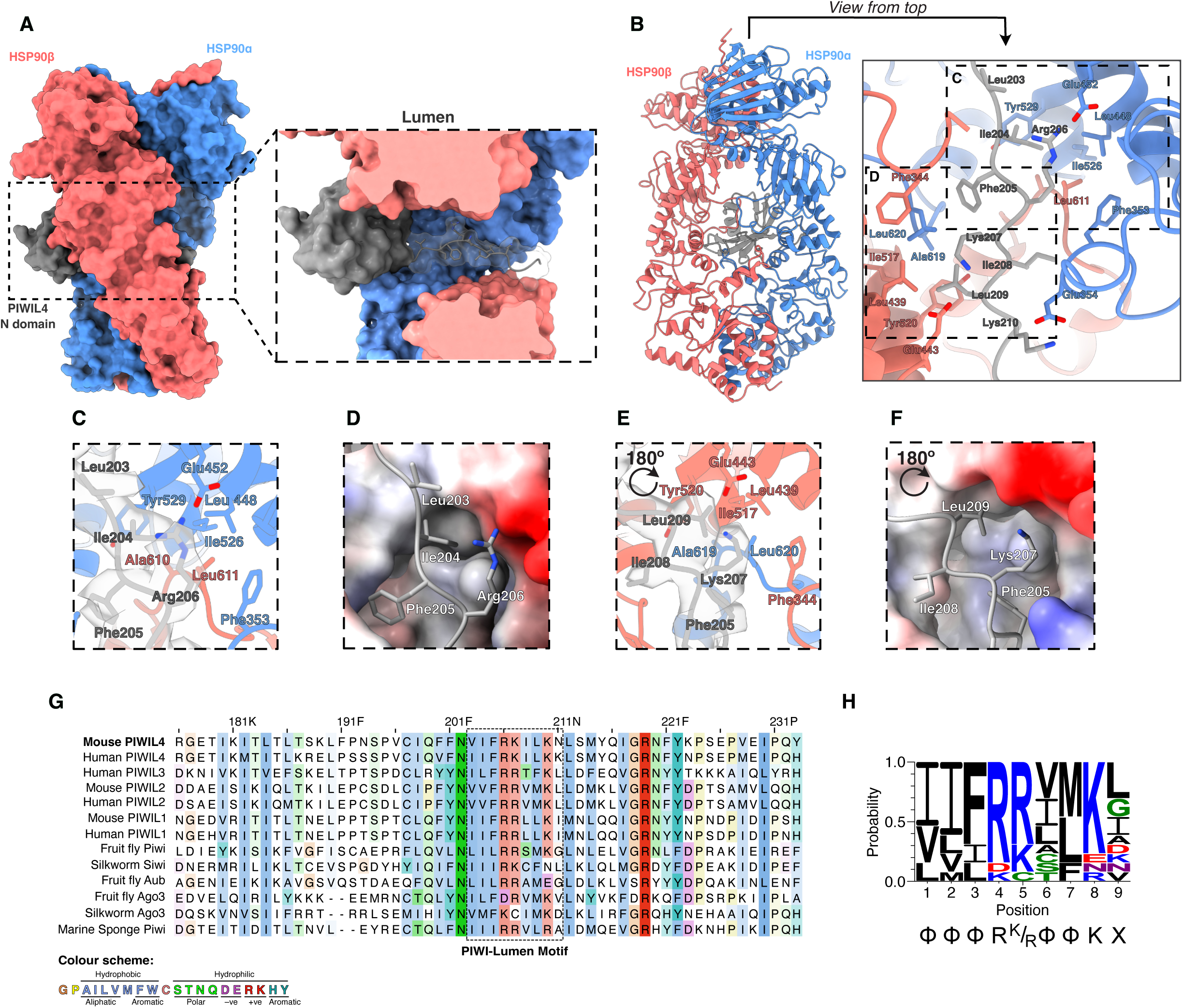
A conserved motif within PIWIL4’s Linker 1 passes through the lumen of the HSP90 dimer. **A**, Transversal view of the HSP90 lumen showcasing the binding of PIWIL4 Linker 1. **B,** Cartoon representation of the resolved regions of the mouse PIWIL4 structure with view from the top of the PIWIL4 and HSP90 residues that drive the lumen interaction. **C,** Close-up view of PIWIL4 Arg206 and neighbouring residues fitted in the cryo-EM density map (only density of the lumen peptide is shown for clarity). **D,** Electrostatic potential of HSP90 surface that surrounds PIWIL4 Arg206 and upstream hydrophobic residues of Linker 1. **E,** Close-up view of PIWIL4 Lys207 and neighbouring residues fitted in the cryo-EM density map (only density of the lumen peptide is shown for clarity). **F,** Electrostatic potential of HSP90 surface that surrounds PIWIL4 Lys207 and downstream hydrophobic residues of Linker 1. **G**, Multiple sequence alignment of part of the Linker 1 region from the indicated PIWI family proteins from the indicated species. Residues with > 20% sequence identity conservation are coloured by physicochemical similarity (Clustal colour scheme from Jalview). Conservation is indicated by colour saturation. **H**, Logo plot summarizing residue frequencies at positions contacting the HSP90 lumen across metazoan PIWI proteins. Below the logo plot is the PIWI-HSP90 consensus lumen motif, X – any residue, φ – hydrophobic residue.

We next examined whether the mode of PIWIL4 L1 engagement with the HSP90 lumen is conserved in other evolutionarily related PIWI proteins. Amino acid sequence conservation analysis of PIWI proteins from organisms across metazoans (Figure 3G) revealed high sequence conservation within the L1 region, with a signature motif characterised by two basic residues flanked by hydrophobic amino acids on either side, followed by an additional basic residue (Figure 3H). The conservation of this motif across PIWI evolution suggests that the mechanism of HSP90-mediated PIWI structural rearrangement is a conserved feature of PIWI proteins.

### HSP90-mediated conformational reconfiguration of PIWI proteins is universal

To test this hypothesis, we next expressed human FLAG-PIWIL2 in ESCs. As with mouse PIWIL4, human PIWIL2 protein co-precipitated HSP90α, HSP90β, HSC70, STIP1, FKBP4, and P23 (Figure 4A). ATP/molybdate treatment also remodelled the respective complexes, with loss of STIP1 and reduced HSC70 binding (Figure 4A). The HSC70 and HSP90 chaperone system is extremely highly conserved^50^. We took advantage of this conservation to determine whether invertebrate PIWI proteins form chaperone complexes in ESCs. Expression and purification of fruit fly Piwi (*Drosophila melanogaster)* or the marine sponge (*Amphimedon queenslandica*) Piwi from ESCs revealed the same associated chaperones (Figure 4A). Furthermore, ATP/molybdate treatment induced the expected co-chaperone remodelling in the fly and sponge Piwi complexes (Figure 4A). To determine whether the HSP90-bound PIWIL4 conformation is conserved across PIWI family members, we determined the cryo-EM structures of ATP/molybdate-treated human PIWIL2 and fruit fly Piwi at 9.8 Å and 3.4 Å resolution (Figure 4B-E, Figure S3 and Figure S4), respectively. Both structures revealed an architecture similar to the mouse PIWIL4 complex, with HSP90 in an ATP-bound closed state and the reconfigured PIWI protein in an extended conformation with the N domain and RNA-binding PAZ/MID/PIWI domains flanking either side of the lumen (Figure 4D-F, also observed in 2D classes in Figure 4B-C). The density of the lumen peptide in the fly structure, with a local resolution ranging between 3.0 and 3.5 Å (Figure S3G), showed distinct side chain features (Figure 4F), which allowed us to model the fly Piwi L1 lumen peptide using mouse PIWIL4 as a template. Consistent with the mouse PIWIL4 structure, the ‘signature motif’ of fly Piwi L1 engages the lumen via a combination of hydrophobic (Leu199, Ile200, Leu201 and Met205) and electrostatic (Arg202, Arg203, Lys207) interactions (Figure 4H-L). Thus, HSP90-mediated conformational opening of PIWI proteins involves the specific recognition of this conserved L1 signature motif by the HSP90 lumen, stabilizing an extended open conformation in which the N domain lies on one side of the cavity and the RNA-binding PAZ/MID/PIWI domains lie on the other (Figure 4M).

**Figure 4:**
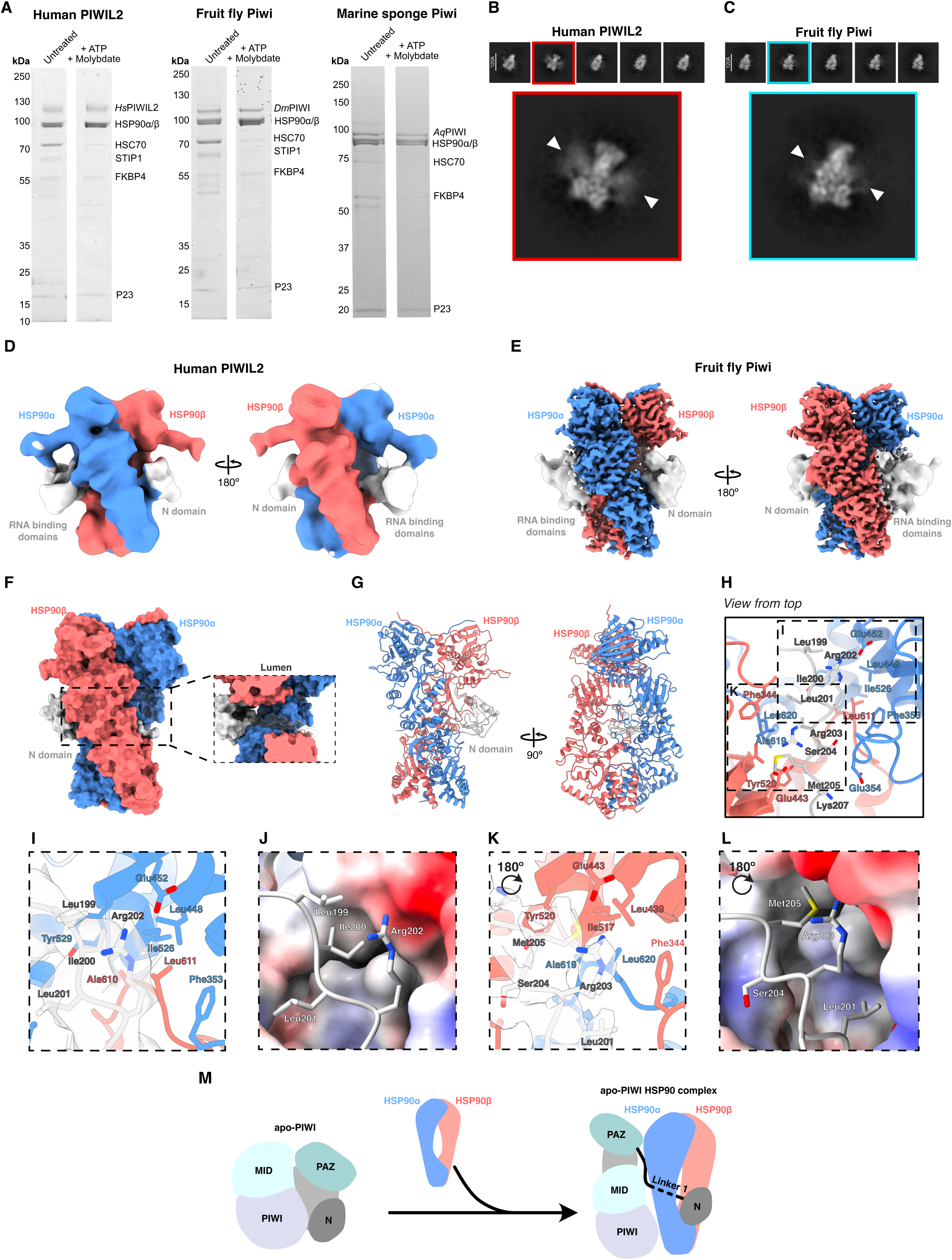
The apo-PIWI-HSP90α/β open conformation and lumen-binding peptide are evolutionarily conserved. **A**, Representative Coomassie-stained SDS-PAGE gel of anti-FLAG immunoprecipitates from ESCs expressing the indicated 3xFLAG-3C-HA or 3xFLAG-3C-HA-tagged human, fruit fly or marine sponge PIWI proteins lysed either in the absence (untreated) or presence (+) of ATP and sodium molybdate. For uncropped whole gel source images, see Supplementary Fig. 1Z. **B,** Representative 2D classes of PIWIL2-HSP90 complex, with an enlarged class showing densities flanking the HSP90 lumen (indicated by white arrows). **C**, same as **B** for fruit fly PIWI. **D**, Cryo-EM density map of human PIWIL2 (grey) in complex with HSP90α (blue) and HSP90β (salmon) in the HSP90 closed state. **E**, Cryo-EM density map of fruit fly Piwi (light grey) in complex with HSP90α (blue) and HSP90β (salmon) in the HSP90 closed state. **F**, Transversal view of the HSP90 lumen showcasing the binding of fruit fly PIWI Linker 1. **G,** Cartoon representation of the resolved regions of the fruit fly PIWI structure in two orientations, in the same colours as described in **E**. **H,** View from the top of the residues in fruit fly Piwi and HSP90 that drive the lumen interaction. **I,** Close-up view of Piwi Arg202 and neighbouring interactions fitted in the cryo-EM density map (only lumen peptide shown). **J,** Surface electrostatic potential of the upstream hydrophobic pocket. **K**, Close-up view of Piwi Arg203 and neighbouring residues fitted in the cryo-EM density map. **L,** Electrostatic surface potential of the downstream hydrophobic pocket in the fly PIWI structure. **M**, Schematic model of PIWI loading onto HSP90 to form the apo-PIWI-HSP90α/β complex.

## Discussion

Here, we show that HSC70, HSP90 and co-chaperones associate with mouse PIWIL2 and PIWIL4 in E16.5 foetal testis. At this stage of development, the piRNA pathway is fully engaged in post-transcriptional transposon silencing, piRNA amplification, and piRNA-directed DNA methylation^1,2,10,46,51,52^. We further demonstrate that mouse PIWIL4 forms complexes with HSC70/HSP90 chaperones and co-chaperones when expressed in ESCs. These cells lack piRNA biogenesis pathways; hence PIWIL4 is in its apo form and engaged with chaperones. The presence of HSC70 and STIP1 suggests that, like the glucocorticoid receptor, these factors load HSP90 onto PIWIL4. The reduction of HSC70 and STIP1 upon ATP/molybdate treatment also supports this model. We also demonstrate that the formation of these complexes is a conserved feature of PIWI proteins from human to the most ancient marine sponge proteins. The structure of apo-PIWIL4 bound to HSP90 reveals a striking reorganization of PIWIL4 into an open conformation. We also demonstrate that the open conformation is a conserved structural feature observed in both human PIWIL2 and fly Piwi proteins. These observations are in agreement with the recently published structure of AGO2-HSP90β-P23 and duplex RNA^44^. In this structure, AGO2 is bound by a HSP90 dimer in a similar manner to PIWI proteins, with the N domain and RNA-binding module oriented on opposite sides of the HSP90 dimer^44^. In our PIWI structures, we resolved the N domain and lumen peptide but not the RNA-binding module, likely due to the lack of either pre-piRNA or FKBP co-chaperones. Indeed, duplex RNA was required to solve the structure of the HSP90-bound AGO2 and its RNA-binding module^44^. We found that like AGO2, HSP90 binds to the L1 domain of PIWI proteins, albeit the site of binding slightly differs. Our structure also differs with regard to the HSP90 variants; the AGO2 complex was resolved with HSP90β homodimers, whereas our PIWI structures were with HSP90α/β heterodimers. This difference could be due to differences in experimental conditions, with the AGO2 study overexpressing AGO2 and HSP90β^44^, whereas we co-purified endogenous HSP90α/β with PIWI proteins. We found that the PIWI peptide bound by the HSP90-lumen was highly conserved across metazoa. This conservation, along with the mouse PIWIL4 and fly Piwi structures allowed us to confidently define a consensus binding motif and provide the structural basis for its binding by the HSP90 dimer. This motif and mechanism of binding are similar to other HSP90-client interactions^41–44,48,49,53,54^ sharing diverse hydrophobic residues common to HSP90 lumen peptides, but also including basic residues that appear unique to PIWI-HSP90 interactions. Based on our PIWI structures and the intriguing AGO2 structures^44^, we propose a speculative model whereby the open PIWI conformation observed in the HSP90 complex is required for pre-piRNA binding (Figure 4M). In summary, here we show that PIWI-HSP90 complexes, the open apo-PIWI conformation, and the HSP90 lumen-binding peptide are conserved characteristics of PIWI proteins.

## Methods

### Mouse strains and experimentation

The *Piwil4^HA^* (*Miwi2^HA^*) mouse allele has been described previously^45^. The *Piwil4^HA^* mice were kept on a mixed B6CBAF1/Crl;C57BL/6N;Hsd:ICR (CD1) genetic background. The *Piwil2^HA^* (*Mili^HA^*) allele was generated by CRISPR-Cas9 gene editing as previously described^46^. A single sgRNA (GCCTCTCAATGGATCCTGTC) together with CAS9 mRNA and a single-stranded DNA oligo containing a GS flexible linker and the 9 amino acid haemagglutinin (HA) epitope tag flanked by 70 nucleotides of left and 70 nucleotides of right homology arm (AATCCTTTGAAAATTCTGGCAGATAAGGAATTGGAACAGAACCAACAGTCGTTTCTACAGCCTCTCAATGggatcctacccatacgatgttccagattacgctGATCCTGTCAGGCCGTTGTTCAGGG GGCCCACCCCAGTCCACCCATCTCAGTGTGTGCGGATGCCAGGCTGTTGGCCTCAAG) was injected into the cytoplasm of fertilised 1-cell zygotes (B6CBAF1/Crl). F_0_ offspring were screened by PCR and the *Piwil2^HA^* allele was confirmed by Sanger sequencing. The allele was established from one founder animal and back-crossed multiple times to a C57BL6/6N genetic background. Thus, the *Piwil2^HA^* mice were on a mixed B6CBAF1/Crl;C57BL/6N genetic background, except for IP-MS experiments where homozygous male studs were mated to Hsd:ICR (CD1) females so that the collected foetuses were on a mixed B6CBAF1/Crl;C57BL/6N;Hsd:ICR (CD1) genetic background. Mice were genotyped by a 2-primer PCR (F – TGCATCTCTCAACTCCCATCAG, R – GCCTTACCTTTTGTACTGGCTG).

Male fertility was assessed by mating studs to C57Bl6/J wild-type females and counting the number of pups born for each plugged female. For each experiment, animal tissue samples were collected from one or more litters and allocated to groups according to genotype. No further randomization or blinding was applied during data acquisition and analysis.

Animals were maintained at the University of Edinburgh, UK, in accordance with the regulation of the United Kingdom’s Home Office. Ethical approval for the UK mouse experimentation has been given by the University of Edinburgh’s Animal Welfare and Ethical Review Body and the work done under license from the United Kingdom’s Home Office. Mice were housed under the following conditions: Lighting: 12-hour light to dark cycle from 7 AM to 7 PM; Temperature: 20-24 °C; Humidity: 45-65 %; Caging: Tecniplast GM 500 Individual Ventilated Cages; Cage substrate: Aspen chips; Enrichment: Aspen chew sticks, carboard dome home, rodent roll and plastic tube for non-aversive handling.

### Immunoprecipitation and mass spectrometry

The anti-HA IP-MS of HA-PIWIL2 from E16.5 *Piwil2^HA/+^* foetal testes was performed as described previously^46^, and the spectra were analysed using FragPipe^55^. Anti-HSP90α IP-MS was performed using anti-HSP90α (PA3-013, Invitrogen) crosslinked to Protein G Dynabeads (2:3 antibody to beads ratio) from 25 wild-type E16.5 foetal testes per replicate as described previously^56^, with control commercial rabbit serum (NS01L, Sigma-Aldrich) crosslinked to Protein G Dynabeads (2:3 serum to beads ratio) used for the control immunoprecipitation. All statistically significant (P < 0.05) enriched (Enrichment > 2-fold) proteins in the anti-PIWIL2 IP-MS are listed in Supplementary Table 1. All statistically significant (P < 0.05) enriched (Enrichment > 2-fold) proteins in the anti-HSP90α IP-MS are shown in Supplementary Table 2.

### Immunofluorescence

Immunofluorescence (IF) experiments were done on freshly cryo-sectioned OCT-embedded E16.5 foetal testis samples (6 μm sections) as previously described^23^. The following primary antibodies were used in this study – rabbit monoclonal anti-HA (clone C29F4, 3724, Cell Signaling Technologies, lot 12) 1:200, rat monoclonal anti-HA (clone 3F10, ROAHAHA, Roche, lot 65506600) 1:200, rabbit polyclonal anti-HSP90α (PA3-013, Invitrogen, lot YI382583) 1:1000, rabbit polyclonal anti-HSP90β (PA3-012, Invitrogen, lot YI375518) 1:1000, mouse monoclonal anti-HSC70 (clone 1A4B3, 66442-1-Ig, ProteinTech, lot 10020187) 1:200. All primary incubations were performed overnight at 4 °C. Secondary antibodies used were – donkey anti-rabbit IgG Alexa Fluor 488 (A-21206, Invitrogen) or 568 (A10042, Invitrogen), donkey anti-mouse IgG Alexa Fluor 568 (A10037, Invitrogen) or 647 (A-31571, Invitrogen), and donkey anti-rat IgG Alexa Fluor 488 (A-21208, Invitrogen), 568 (A78946, Invitrogen) or 647 (A48272, Invitrogen), all at a dilution of 1:1,000 in blocking buffer with 10 μg/ml DAPI. All secondary incubations were for 1 hour at room temperature.

Images were taken on a Zeiss LSM980 with Airyscan II module in Super-Resolution (SR) mode, with optimal pinhole settings for SR imaging. The most restrictive excitation and emission filters available were chosen to prevent signal bleed-through between channels. Images were SR-processed with the Zeiss Zen 3.5 software “Airyscan processing” function with settings “3D” and a strength of “Auto”.

### Cell culture and transfection

Mouse embryonic stem cells (E14Tg2a, a kind gift from Prof. Keisuke Kaji, University of Edinburgh) were cultured and transfected as described previously^57^. Cells expressing 3xFLAG-tagged PIWI proteins were obtained by random integration of a protein-of-interest-IRES-GFP cassette by a *PiggyBac* transposase system as described previously^57^. Cells were sorted for high GFP expressing cells 5 days and 8 days post transfection. All cells were regularly checked for mycoplasma contamination.

### Protein purification

PIWI-overexpressing ESC pellets were thawed and lysed in 10 mL of lysis buffer (Buffer A (20mM HEPES-pH 7.5, 100mM Potassium Acetate-pH 7.5, 3.5mM Magnesium Acetate-pH 7.5, 1mM DTT) + 5% Glycerol, cOmplete ULTRA EDTA-free protease inhibitor, 0.1% v/v Triton X-100) for every gram of cell pellet. ATP/molybdate conditions contained an additional 20 mM sodium molybdate and 5 mM ATP in all buffers. The lysate was incubated on ice for 30 minutes and the lysate was then clarified by centrifugation at >21,000 RCF for 30 min at 4 °C. Anti-FLAG G1 resin (L00432S, Genscript) was prepared by equilibrating 10 μL (1CV) of packed resin per 1 mL of clarified lysate with 100 column volumes (CV) of lysis buffer, repeated twice, followed by blocking with 200 CV of blocking buffer (Buffer A + 5mg/ml BSA) for 1 hour at 4 °C. The resin was washed once with 10CV of lysis buffer and the clarified lysate was applied and incubated with rotation for 2h at 4 °C. The resin was subsequently washed twice with 20CV of Buffer A + 5% Glycerol and once with 20 CV of Buffer A only. Bound PIWI protein was eluted using 5 CV of elution buffer (Buffer A + 500 ng/ul 3xFLAG-Peptide (RP21087-10, Genscript)) on ice for 1 h. Finally, the eluate was concentrated using a 30kDa Vivacon 500 concentrator spin column (VN01H22, Sartorius) until the desired protein concentration was reached. Due to the presence of the FLAG peptide in the eluted sample, the accurate concentration of the FLAG-purified complexes could not be measured by traditional spectrophotometric methods.

### Cryo-EM sample preparation and data collection

FLAG-purified ATP/molybdate-treated samples containing mouse PIWIL4, human PIWIL2 and fly PIWI were crosslinked with 0.01 % (v/v) glutaraldehyde for 5 min at 4 °C and quenched with 50 mM Tris-HCl pH 7.5. The three PIWI-containing complexes were applied onto Quantifoil R2/1 300-mesh grids coated with 2 nm of carbon previously glow-discharged for 7 s with a current of 20 mA in the presence of air. 4.5 μl of sample were added to the grids and blotted for 2.5 s at 10 °C and 95% humidity using a Leica EM GP plunger (Leica). Prior to vitrification, 5 mM NDSB-256 and 0.02 % Glyco-diosgenin were added.

Data was collected in a Titan Krios transmission electron microscope operating at 300 keV, equipped with a Falcon 4i direct electron detector and a Selectris X Energy Filter with an energy slit width of 5 eV (ThermoFisher). Data was acquired at a nominal magnification of x165,000, which resulted in a pixel size of 0.727 Å/px. Exposure time was adjusted to keep a constant total dose of 40 e/ Å^2^ per movie, with a defocus range varying between −2.6 μm and −0.5 μm in steps of 0.3 μm. Grid screening and automated data acquisition was performed with Smart EPU software (ThermoFisher). Parameters used for each collection are summarized in Supplementary Table 3.

### Cryo-EM data processing and model building

All datasets were processed with CryoSPARC v4.7 and v5. ^58^ (Figure S2-4)

For mouse PIWIL4, 63,060 movies were collected. Following patch motion correction and CTF estimation, an initial subset of 2,101 denoised micrographs was used to manually pick 304 particles that were subsequently used for template-picking. After a few rounds of 2D classification, 23,570 particles that yielded classes showcasing secondary structure features were used to train a Topaz picking model^59^ that was then applied to the entire dataset. After iterative rounds of 2D classification combined with Topaz training, 74,944 particles were subjected to ab initio reconstruction with two classes followed by heterogeneous refinement. Homogeneous refinement of the best heterogeneous refinement class yielded an initial model at 6.12 Å resolution containing 46,324 particles, used to train Topaz. Recursive rounds of Topaz picking followed by 2D classification resulted in 444,368 particles with corresponding 2D classes displaying high resolution features. Up to this point, processing was performed with particles extracted with a box size of 400 px binned to 100 px (2.9 Å/px).

Following re-extraction with full box size (0.727 Å/px), multiple rounds of heterogeneous refinement using the initial 6.12 Å model and junk classes as inputs were performed to isolate a high-quality pool of 175,531 particles. Non-uniform refinement (NU-refinement) of this pool produced a 3.04 Å reconstruction. Focused 3D classification with a mask around the PAZ/MID/PIWI domains isolated a class with 114,414 particles that yielded a 3.18 Å model via non-uniform refinement. After global and local CTF refinement, a final round of NU-refinement was performed on the remaining 106,227 particles, producing the final HSP90-lumen map at a global resolution of 3.16 Å (FSC = 0.143). This same particle stack was also subjected to local refinement using a focus mask centered on the N domain. The HSP90-lumen map (for the PAZ/MID/PIWI domains) and the locally refined map (N domain) were locally filtered in CryoSPARC based on local resolution estimation; and a composite map was generated in ChimeraX combining the HSP90-lumen map with both locally filtered maps.

For fly PIWI, 26,580 movies were collected and patch motion correction and CTF estimation was performed. A mouse PIWIL4 Topaz model was used to pick an initial subset of 954,463 particles extracted with a binned box size of 100 px (pixel size of 2.9 Å/px). After several rounds of 2D classification, 750,378 particles yielding 2D classes with secondary structure features were selected. These particles were used in two parallel processing branches. In the first branch, to isolate particles containing HSP90s in the closed state, heterogeneous refinement was performed recursively, using a mouse PIWIL4 model and several junk classes as inputs. This resulted in a subset of 141,884 particles that produced a 3.64 Å reconstruction by NU-refinement after re-extraction with a full box size (400 px, 0.727 Å/px). A round of ab initio reconstruction with two classes, followed by heterogeneous refinement, resulted in a map of fly PIWI bound to HSP90 containing 106,459 particles.

Parallel to this, 33,950 particles within the 750,378 particles pool were used to train Topaz. Particles were extracted with a box size of 400 px (0.727 Å/px) and subjected to multiple rounds of 2D classification. 409,312 good particles were further cleaned up by heterogeneous refinement using mouse PIWIL4 as one of the templates. The resulting 157,737 particles (which produced a 3.42 Å reconstruction by NU-refinement), were further classified by ab initio reconstruction followed by heterogeneous refinement. Particles corresponding to one of the classes (86,358 particles) were pooled with the 106,459 particles of the first processing branch and cleaned up by subsequent ab initio reconstruction followed by heterogeneous refinement. After NU-refinement, a final map of HSP90 with occupied lumen at 3.35 Å was obtained. Focused 3D classification with a mask centered around the N domain followed by NU-refinement resulted in a map with medium-resolution features around the N domain. A composite map was generated by combining the final HSP90-lumen map with a locally filtered map of the region corresponding to the PAZ/MID/PIWI-domain and the NU-refinement map with resolved N-domain.

For human PIWIL2, 28,170 movies were recorded. After patch motion correction and CTF estimation, the same mouse PIWIL4-trained Topaz model was used to pick particles (box size 100 px, 2.9 Å/px). After several rounds of 2D classification, 23,857 particles were used to generate a Topaz picking model which. After extraction with full box size (400 px, 0.727 Å/px) 2D classification, 245,459 particles were cleaned up using heterogeneous refinement with mouse PIWIL4 as one of the inputs. Resulting 45,324 particles were further classified by ab initio followed by heterogeneous refinement, with the best class refined by NU-refinement. A final map with a resolution of 9.83 Å was obtained.

For model building, a model of the HSP90α/β heterodimer was generated with AlphaFold3 ^60^ and rigid-body fitted in the cryo-EM maps using in ChimeraX v1.12. Mouse PIWIL4 lumen peptide was modelled by amino acid substitution using the structure of Ago2 bound to HSP90 (PDB 9W5I) as a template. Fly PIWI lumen peptide was modelled using mouse PIWIL4 as a template. Models were subjected to iterative rounds of real-space refinement using Phenix (v1.21.2), Coot-1 and Namdinator^61^. Models for the N domains were predicted by AlphaFold3 and fitted against the corresponding maps with Namdinator.

### Protein sequence conservation analysis and motif prediction

Multiple sequence alignments for PIWI proteins was generated with Clustal Omega ^49^ and visualised in Jalview (2.11.4.1) ^50^ for the following canonical sequences from UniProt ^51^ – Mouse (*Mus musculus*) PIWIL4 (Q8CGT6), Mouse (*Mus musculus*) PIWIL2 (Q8CDG1), Mouse (*Mus musculus*) PIWIL1 (Q9JMB7), Human (*Homo sapiens*) PIWIL4 (Q7Z3Z4), Human (*Homo sapiens*) PIWIL3 (Q7Z3Z3), Human (*Homo sapiens*) PIWIL2 (Q8TC59), Human (*Homo sapiens*) PIWIL1 (Q96J94), Fruit fly (*Drosophila melanogaster*) Piwi (Q9VKM1), Fruit fly (*Drosophila melanogaster*) Aub (O76922), Fruit fly (*Drosophila melanogaster*) Ago3 (Q7PLK0), Silkworm (*Bombyx mori*) Siwi (A8D8P8), Silkworm (*Bombyx mori*) Siwi (A9ZSZ2), Marine sponge (*Amphimedon queenslandica*) Piwi (A0AAN0JJ68). Lumen peptide motif logo was visualised using WebLogo 3.9.0 ^62^.

### Foetal testes extract preparation for western blotting

Foetal testis protein extracts for western blot analysis were prepared and analysed as described previously^46^. Blots were probed overnight at 4 °C with rabbit anti-HA (3724, Cell Signaling Technologies, 1:1,000) and mouse anti-alpha tubulin (T9026, Sigma, 1:1,000) in blocking buffer (5% skim milk in TBS with 0.1% Tween 20). IRDye 680RD donkey anti-mouse IgG (926-68072, LICOR Biosciences) and IRDye 800CW donkey anti-rabbit IgG (926-32213, LICOR Biosciences) secondary antibodies were used at 1:20,000 in blocking buffer at room temperature for 1 hour, and the blots were imaged on a Chemidoc MP imager (Bio-Rad). Exposure of the whole blot was adjusted for presentation with ImageJ.

## Quantification and statistical analysis

Data were plotted in R (version 4.4.2 (2024-06-14)) using the ggplot2, tidyr, dplyr, ggpubr and Hmisc toolkits (versions ggplot2_3.5.1, tidyr_1.3.1, dplyr_1.1.4, ggpubr_0.6.0, Hmisc_5.2.1), Python (version 3.12.9) using the pandas, scipy, scikit-learn, matplotlib and seaborn packages (versions pandas_2.2.3, scipy_1.14.1, scikit-learn_1.5.2, matplotlib_3.9.2, seaborn_0.13.2) or Microsoft Excel for Mac (Office 365, version 16.9). For the IP-MS data, statistical testing was performed with R (version 4.4.2 (2024-06-14)) using the R Studio software and with Perseus (version 1.6.5.0) ^53^. Unpaired, two-tailed Student’s t-tests were used to compare differences between groups and adjusted for multiple testing using Benjamini–Hochberg correction where indicated. Averaged data are presented as mean ± SEM (standard error of the mean) for comparisons or mean ± SD (standard deviation) for descriptive statistics, unless otherwise indicated. No statistical methods were used to predetermine sample size. The experiments were not randomized, and the investigators were not blinded to allocation during experiments and outcome assessment unless otherwise noted.

## Data availability

Data for the IP-MS experiments were deposited at ProteomeXchange under the accession number PXDXXXXXX. HA-PIWIL4 IP-MS data was re-analysed from ProteomeXchange accession PXD016701. PDB and EMDB accession codes are 32BJ and EMD-58778 for mouse PIWIL4; 32BM and EMD-58779 for fly PIWI; EMD-58780 for human PIWIL2 (Supplementary table 3).

## Acknowledgements

This research was supported by the Wellcome Trust funding to D.O.C. (225237), A.G.C. (200898), A.A.J (309153/Z/24/Z), the Wellcome Centre for Cell Biology (203149) and the multi-user equipment grants (108504 and 092076). A.A.J. and his team are co-funded by the European Union (ERC, CHROMSEG, grant no. 101054950). The views and opinions expressed are, however, those of the authors only and do not necessarily reflect those of the European Union or the European Research Council. Neither the European Union nor the granting authority can be held responsible for them. This work was supported by funding for the Wellcome Discovery Research Platform for Hidden Cell Biology (226791), and we gratefully acknowledge support from the Microscopy, Proteomics, and Bioinformatics cores. P.J.D. is funded by the Wellcome Trust. D.M.M. was funded by the Wellcome Trust. T.C. was funded by the Darwin Trust of Edinburgh. A.Z. was funded by a German Research Foundation fellowship (DFG, award ZO 376/1-1). This work utilised the Gene Center cryoEM facility funded by LMU Munich and DFG (Deutsches Forschung Gemeinschaft), the University of Edinburgh Protein Production Facility (EPPF), the Wellcome Centre for Cell Biology Centre Optical Instrumentation Laboratory (COIL) / Light Microscopy Core (LMC), Proteomics and Bioinformatics Core Platforms as well as the Centre for Regenerative Medicine’s and the University of Edinburgh School of Biological Sciences FACS facilities. We also thank Karl-Peter Hopfner and his lab members for sharing resources and useful discussions.

## Author contributions

D.M.M. generated the *Piwil2^HA^* mouse allele. D.M.M. identified and isolated the PIWI chaperone complexes. P.D. and P.S.P. performed most biochemical experiments. P.S.P. performed the cryo-electron microscopy and analysis. K.L. and A.R. also performed biochemical experiments. T.C. performed immunofluorescence experiments. A.Z. and D.M.M. performed the IP-MS experiments. C.S. analysed the IP-MS data. D.O’C., A.G.C., and A.A.J supervised this study. D.O’C. conceived this study. P.S.P., A.A.J, D.O’C., P.D., D.M.M. and T.C. wrote the manuscript.

## Resource identification initiative

**Supplementary Figure 1:**
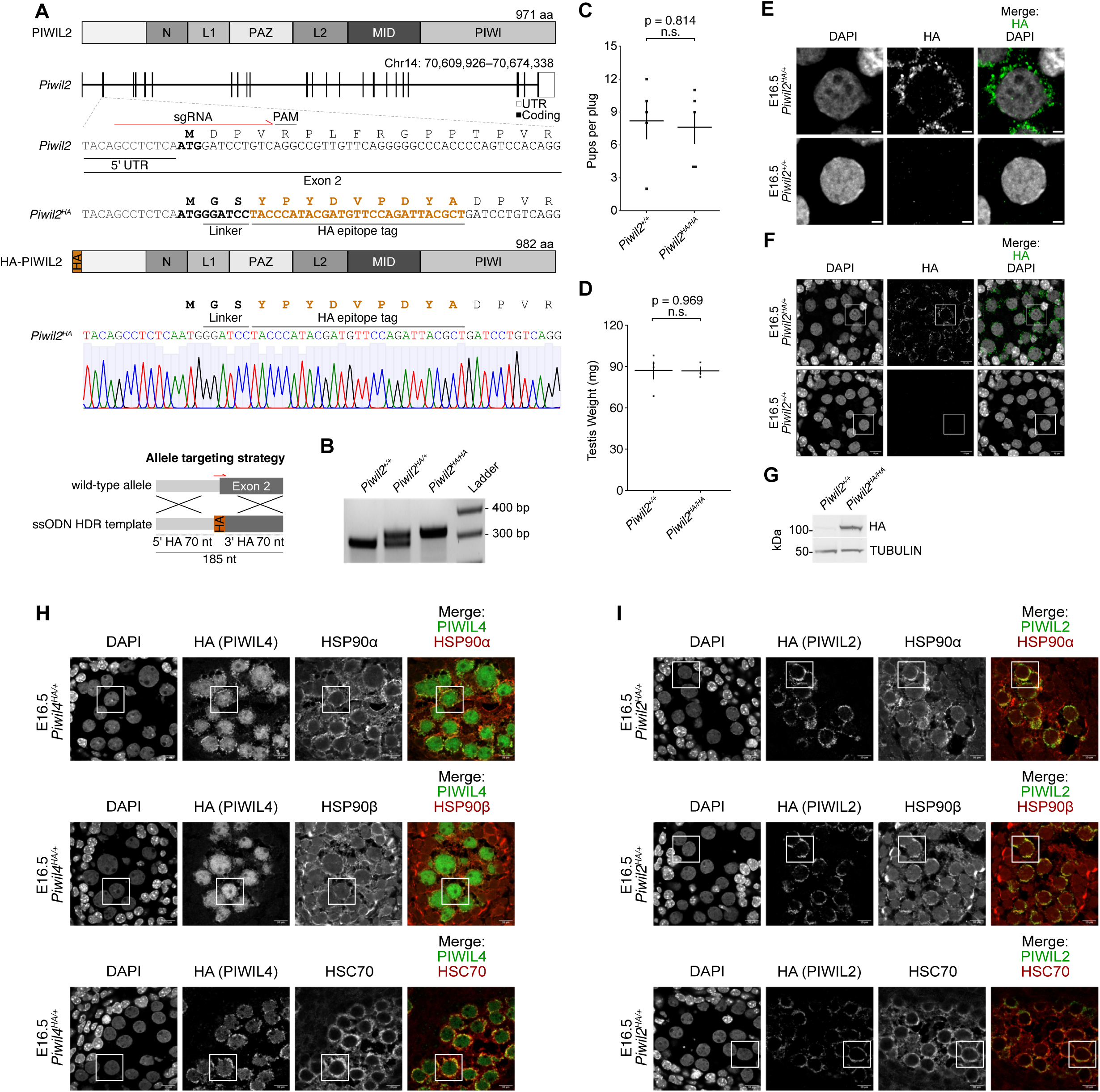
Generation and validation of the Piwil2HA mouse allele. **A**, Schematic representations of the mouse *Piwil2* locus and encoded 971 amino acid protein are shown, along with a schematic of the CRISPR targeting strategy showing the location of single-stranded oligo DNA donor (ssODN) and homology arms (HA) used. The sgRNA used for generation of the *Piwil2^HA^* allele (red) and adjacent PAM sites are indicated, along with a representative sequencing trace of *Piwil2^HA^* exon 2 harbouring the 33-nucleotide insertion encoding a flexible linker and the HA (haemagglutinin) epitope tag (highlighted in brown). Sequencing was performed on n = 1 F_0_ animal. **B**, Representative image of PCR genotyping result for *Piwil2^+/+^*, *Piwil2^HA/+^* and *Piwil2^HA/HA^* mice. PCR genotyping was performed for over 300 mice with similar results. **C**, Number of pups born per plug (mean ± SEM) fathered by n = 5 wild-type and n = 5 *Piwil2^HA/HA^* studs mated to wild-type C57Bl6/J females. **D**, Testis weight (mean ± SEM) of n = 3 wild-type and n = 3 *Piwil2^HA/HA^* 14-week-old mice. **E**, Representative E16.5 gonocyte stained for HA (HA-PIWIL2) and DAPI in foetal testis sections from *Piwil4^HA/+^* (top row) or wild-type (bottom row) mice. **F**, Representative E16.5 seminiferous cords stained for HA (HA-PIWIL2) and DAPI in foetal testis sections from *Piwil2^HA/+^* (top row) or wild-type (bottom row) mice. White rectangles indicate germ cells shown zoomed-in in (**E**). Images in **E-F** are representative of n = 3 biological replicates. Scale bars, 2 μm (**E**) and 10 μm (**F**). **G**, Representative western blot of n = 3 E16.5 foetal testis samples of the indicated genotypes indicating HA-PIWIL4 protein levels in a single foetal testis. Alpha-tubulin served as loading control. For uncropped whole blot source images, see Supplementary Fig. 1X. **H-I**, Representative E16.5 seminiferous cords stained for HA (HA-PIWIL4 in (**H**) and HA-PIWIL2 in (**I**)), HSP90α (top row), HSP90β (middle row) and HSC70 (bottom row) and DAPI in foetal testis sections from *Piwil4^HA/+^* (**H**) or *Piwil2^HA/+^* (**I**) mice. White rectangles indicate germ cells shown zoomed-in in Figure 1E-F. Images are representative of n = 3 biological replicates. Scale bars, 10 μm.

**Supplementary Figure 2.**
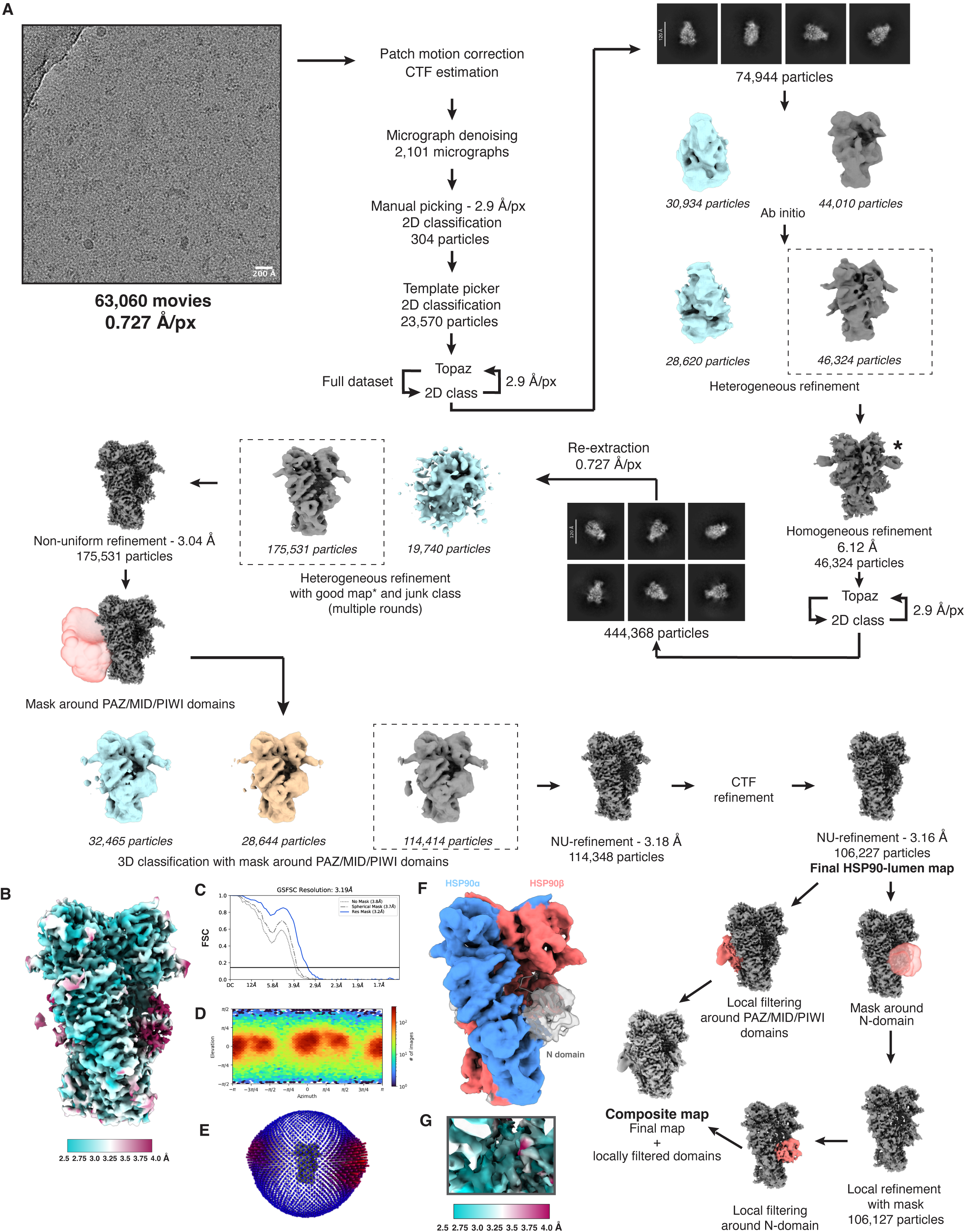
Cryo-EM processing workflow of mouse PIWIL4 and map validation plots. **A,** Cryo-EM data processing workflow. **B,** Local resolution estimation, **C,** FSC curve, **D,** Azimuth plot and **E,** angular distribution of the final HSP90-lumen map of PIWIL4. **F**, Model of the N domain of mouse PIWIL4 fitted in the locally refined map. **G**, Local resolution estimation of mouse PIWIL4 lumen peptide

**Supplementary Figure 3.**
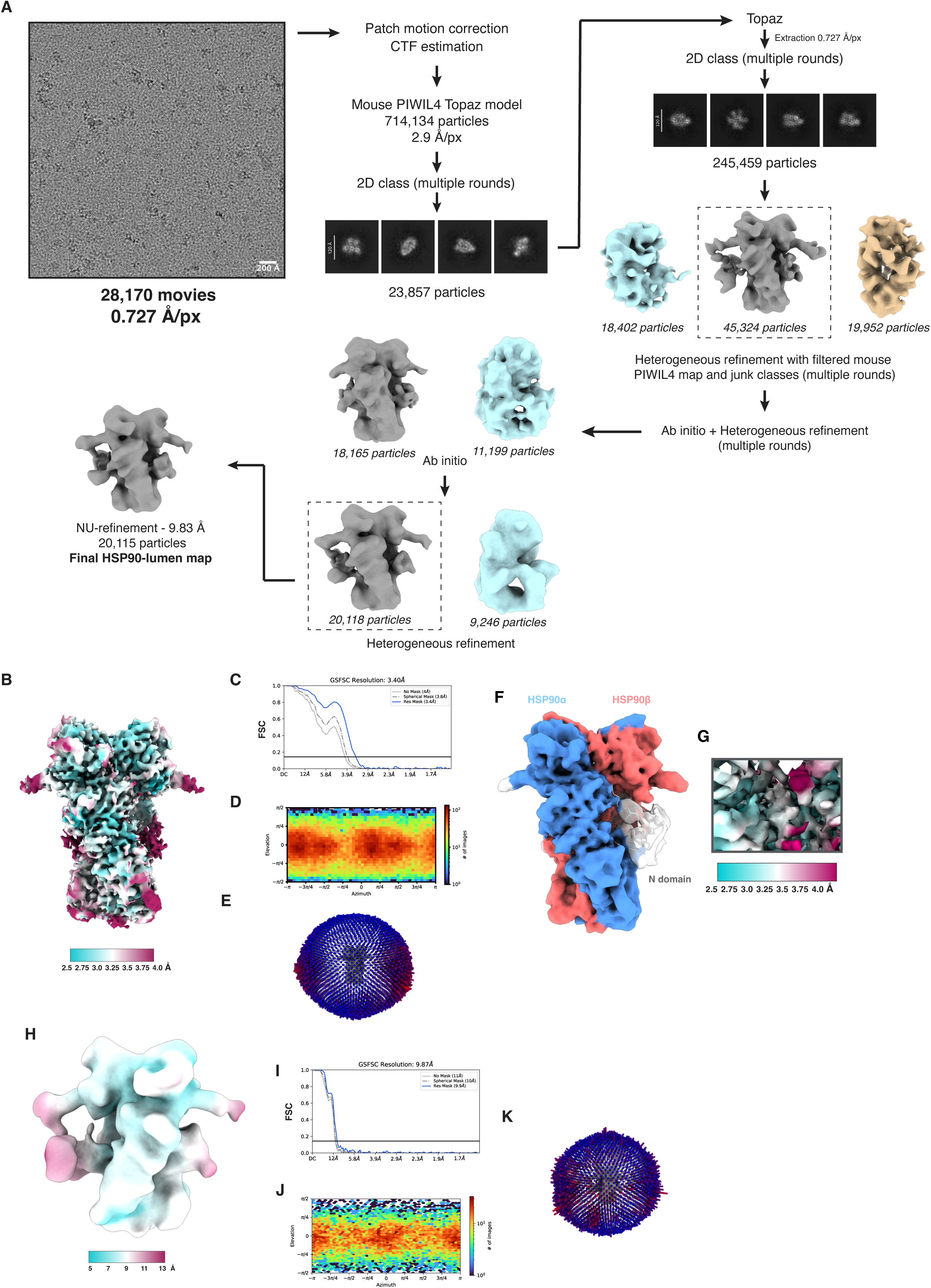
Cryo-EM processing workflow of human PIWIL2 and map validation plots for fly PIWI and human PIWIL2. **A,** Cryo-EM data processing workflow of human PIWIL2. **B,** Local resolution estimation**, C,** FSC curve**, D,** Azimuth plot and **E,** angular distribution of the final HSP90-lumen map of fly Piwi. **F,** Model of the N domain of fly PIWI in the locally refined map**. G**, Local resolution estimation of fruit fly Piwi lumen peptide. **H**, Local resolution estimation **I,** FSC curve**, J** Azimuth plot and **K,** angular distribution of the final human PIWIL2-HSP90 map.

**Supplementary Figure 4.**
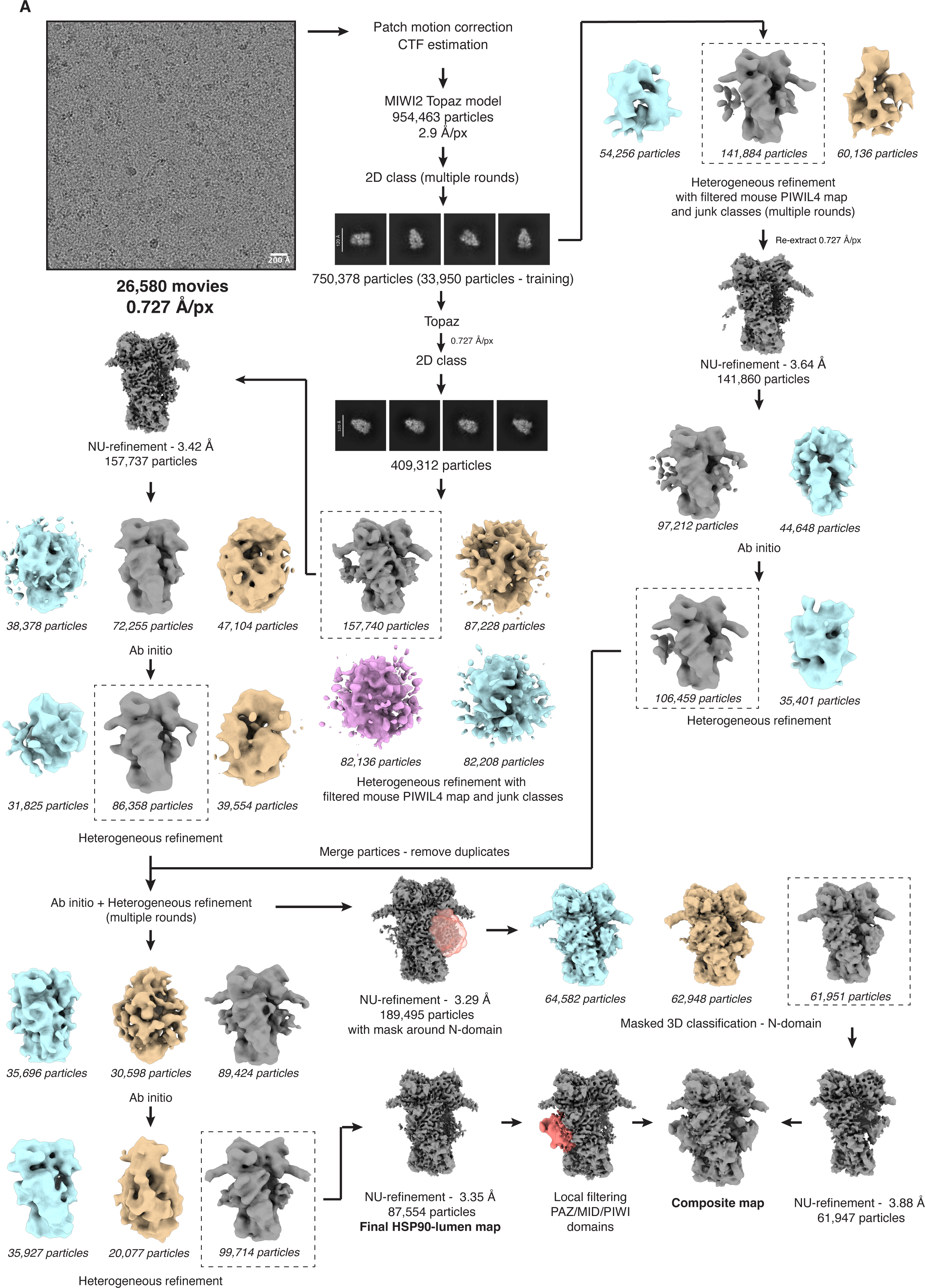
Cryo-EM processing workflow of fruit fly PIWI

**Supplementary Table 1.**
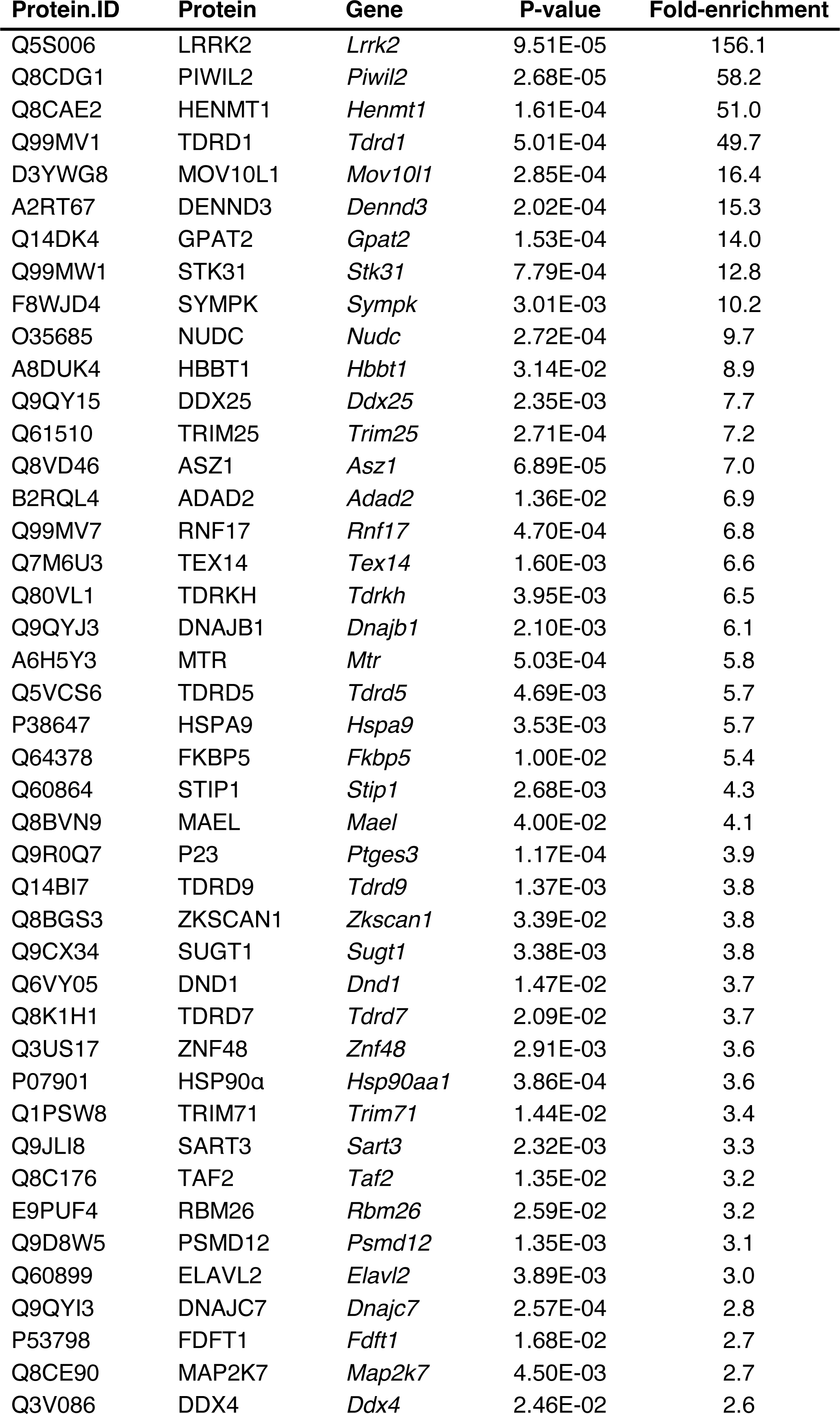
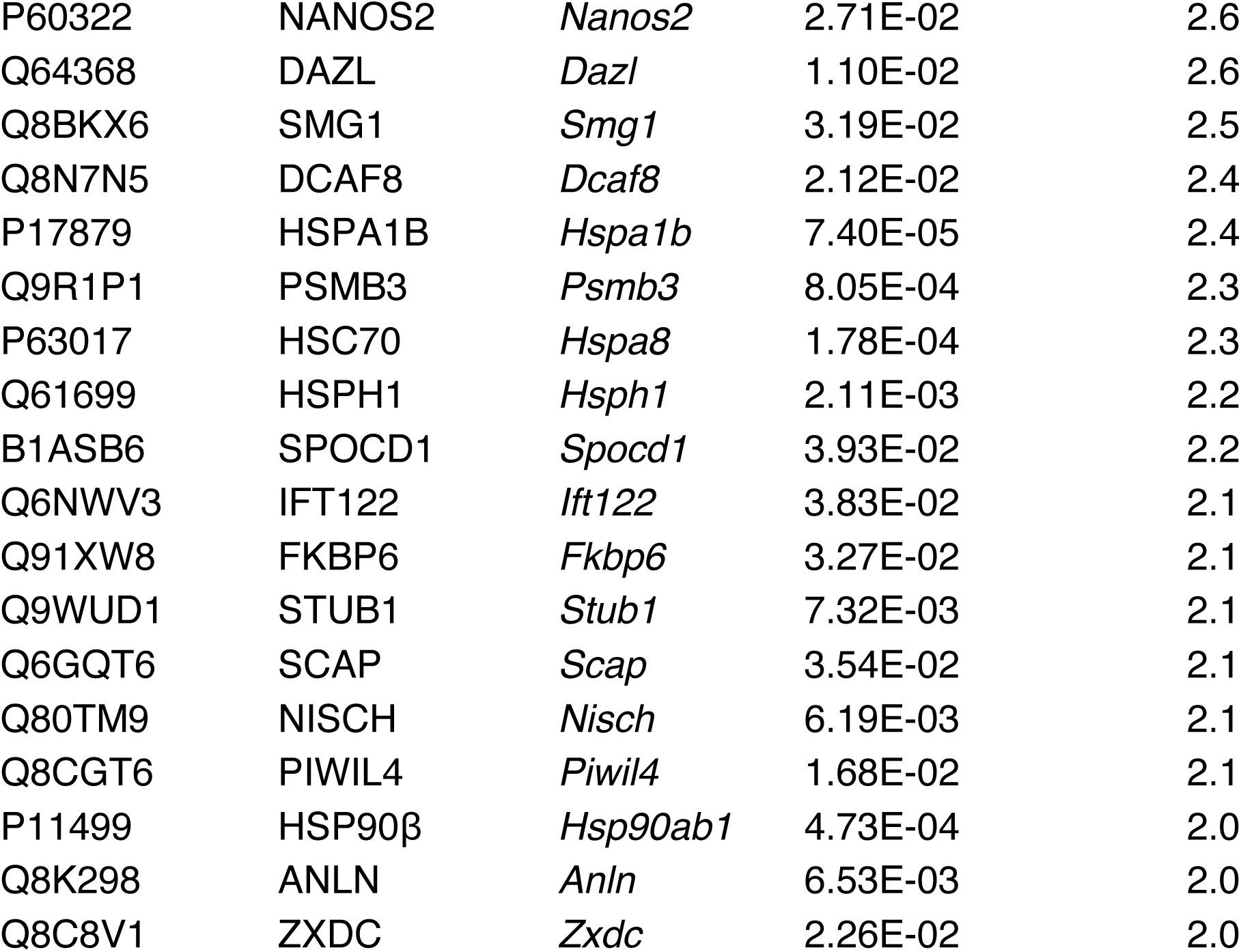
Proteins identified as E16.5 PIWIL2 associations. Table lists all statistically significant, > 2-fold enriched associations identified after immunoprecipitation-coupled mass spectrometry of HA-PIWIL2. Statistical significance threshold was set to P < 0.05 (two-tailed paired Student’s t-test).

**Supplementary Table 2.** Proteins identified as E16.5 HSP90α associations. Table lists all statistically significant, > 2-fold enriched associations identified after immunoprecipitation-coupled mass spectrometry of HSP90α. Statistically significance threshold was set to P < 0.05 (two-tailed paired Student’s t-test).

| Protein.ID | Protein | Gene | P-value | Fold-enrichment |
| --- | --- | --- | --- | --- |
| Q80Y52 | HSP90α | *Hsp90aa1* | 7.0E-125 | 10013.1 |
| Q3THQ5 | STIP1 | *Stip1* | 1.5E-41 | 4305.8 |
| Q71LX8 | HSP90β | *Hsp90ab1* | 8.1E-131 | 4198.9 |
| B1ARU4 | MACF1 | *Macf1* | 3.4E-08 | 1157.8 |
| Q61081 | CDC37 | *Cdc37* | 1.2E-26 | 1127.1 |
| Q9WUD1 | STUB1 | *Stub1* | 6.3E-25 | 513.0 |
| A0A0A6YWX1 | USP19 | *Usp19* | 2.9E-19 | 471.4 |
| Q9CYG7 | TOMM34 | *Tomm34* | 7.9E-30 | 305.0 |
| Q9QYI3 | DNAJC7 | *Dnajc7* | 7.4E-20 | 261.4 |
| Q4FJN2 | FKBP5 | *Fkbp5* | 4.7E-21 | 248.4 |
| Q60676 | PPP5C | *Ppp5c* | 6.9E-28 | 230.6 |
| P17879 | HSPA1B | *Hspa1b* | 4.4E-09 | 206.6 |
| Q8CI32 | BAG5 | *Bag5* | 1.2E-34 | 198.9 |
| P17156 | HSPA2 | *Hspa2* | 4.6E-09 | 198.6 |
| P30416 | FKBP4 | *Fkbp4* | 6.2E-13 | 185.7 |
| Q14BI7 | TDRD9 | *Tdrd9* | 3.3E-07 | 163.7 |
| Q545C3 | CDK4 | *Cdk4* | 2.8E-09 | 158.4 |
| Q9CQJ2 | PIH1D1 | *Pih1d1* | 5.3E-06 | 148.4 |
| P04627 | ARAF | *Araf* | 3.4E-11 | 127.4 |
| Q3TL79 | AHSA1 | *Ahsa1* | 1.4E-15 | 117.5 |
| Q6PB51 | CCDC117 | *Ccdc117* | 3.0E-27 | 113.6 |
| Q9D706 | RPAP3 | *Rpap3* | 3.0E-31 | 109.6 |
| P03976 | KV2A5 | *Kv2a5* | 2.6E-10 | 93.6 |
| A6H5Y3 | MTR | *Mtr* | 4.3E-38 | 77.1 |
| Q8BGF3 | WDR92 | *Wdr92* | 4.1E-07 | 76.4 |
| Q5DTZ0 | NYNRIN | *Nynrin* | 1.6E-07 | 75.8 |
| Q3TJG6 | P23 | *Ptges3* | 3.3E-07 | 73.0 |
| Q9CX34 | SUGT1 | *Sugt1* | 2.7E-21 | 72.2 |
| Q9CZP7 | CDC37L1 | *Cdc37l1* | 1.4E-22 | 70.4 |
| Q9D832 | DNAJB4 | *Dnajb4* | 2.8E-11 | 61.4 |
| Q14DK4 | GPAT2 | *Gpat2* | 3.5E-12 | 60.1 |
| P63017 | HSC70 | *Hspa8* | 5.2E-49 | 57.4 |
| F2YMC1 | MOV10L1 | *Mov10l1* | 6.3E-10 | 51.0 |
| Q7M6U3 | TEX14 | *Tex14* | 6.3E-44 | 50.6 |
| A0A0U1RPG9 | URI1 | *Uri1* | 5.4E-10 | 42.8 |
| E9Q0U7 | HSPH1 | *Hsph1* | 1.7E-08 | 40.3 |
| A2NW55 | E8VHC | *E8vhc* | 1.4E-03 | 40.3 |
| O70591 | PFDN2 | *Pfdn2* | 7.0E-22 | 40.2 |
| Q80YC2 | HSP90AB1 | *Hsp90ab1* | 2.1E-07 | 35.7 |
| Q8C3I8 | HGH1 | *Hgh1* | 9.7E-11 | 35.2 |
| Q3TU79 | DNAJB1 | *Dnajb1* | 6.5E-07 | 34.4 |
| E9Q9G8 | IFT122 | *Ift122* | 8.2E-13 | 32.3 |
| O08915 | AIP | *Aip* | 3.1E-07 | 31.3 |
| Q91YN9 | BAG2 | *Bag2* | 2.1E-23 | 31.2 |
| O35685 | NUDC | *Nudc* | 9.2E-08 | 31.2 |
| Q3UZC4 | TTC4 | *Ttc4* | 1.2E-15 | 30.6 |
| Q8CEJ2 | UXT | *Uxt* | 9.5E-11 | 29.6 |
| X5J4V9 | IGHV3-1 | *Ighv3-1* | 8.2E-08 | 26.8 |
| B0FTY3 | NUDCD3 | *Nudcd3* | 7.5E-11 | 26.6 |
| E9Q9F9 | EDRF1 | *Edrf1* | 2.4E-06 | 26.3 |
| Q8CGT6 | PIWIL4 | *Piwil4* | 2.9E-14 | 25.5 |
| A0A075B5W2 | IGHV1-52 | *Ighv1-52* | 1.1E-11 | 24.0 |
| B2RQL4 | ADAD2 | *Adad2* | 1.0E-08 | 22.3 |
| S4R1Q8 | PLCE1 | *Plce1* | 9.0E-14 | 21.1 |
| Q1W5W7 | GAG-POL | *Gag-pol* | 5.4E-11 | 20.5 |
| Q3TSX8 | TOMM70A | *Tomm70a* | 3.8E-08 | 19.5 |
| Q99N57 | RAF1 | *Raf1* | 3.0E-11 | 18.0 |
| P40336 | VPS26A | *Vps26a* | 1.5E-12 | 17.0 |
| Q99KD5 | UNC45A | *Unc45a* | 8.8E-08 | 16.5 |
| Q8K0T2 | DYNC2LI1 | *Dync2li1* | 1.3E-09 | 16.4 |
| A0A0B4J1K5 | IGLV3 | *Iglv3* | 1.7E-10 | 15.7 |
| Q9CXW3 | CACYBP | *Cacybp* | 6.0E-08 | 15.1 |
| Q3UKN2 | PDRG1 | *Pdrg1* | 2.6E-08 | 14.9 |
| Q66L71 | DIP2A | *Dip2a* | 1.6E-13 | 14.3 |
| A0A0G2JFB2 | TDRKH | *Tdrkh* | 5.2E-26 | 13.4 |
| A0A0G2JER9 | DNAJB6 | *Dnajb6* | 2.6E-09 | 12.7 |
| Q3TRJ1 | VPS35 | *Vps35* | 2.9E-07 | 12.6 |
| Q3TXT7 | RUVBL2 | *Ruvbl2* | 3.9E-39 | 12.0 |
| Q3U1C2 | RUVBL1 | *Ruvbl1* | 1.4E-60 | 10.9 |
| Q3V214 | POLR2E | *Polr2e* | 9.8E-06 | 10.4 |
| P68369 | TUBA1A | *Tuba1a* | 5.8E-05 | 10.4 |
| P26151 | FCGR1 | *Fcgr1* | 9.5E-06 | 10.1 |
| Q99J95 | CDK9 | *Cdk9* | 4.7E-11 | 10.0 |
| Q3U5D9 | AHR | *Ahr* | 3.4E-09 | 9.6 |
| Q80TM9 | NISCH | *Nisch* | 1.4E-06 | 9.6 |
| Q91V83 | TTI1 | *Tti1* | 3.1E-09 | 9.5 |
| Q3TLR3 | TRIM32 | *Trim32* | 3.3E-04 | 9.4 |
| Q9WVL5 | MORC1 | *Morc1* | 1.9E-07 | 9.2 |
| F8WJB0 | MED23 | *Med23* | 2.5E-08 | 9.0 |
| A2AH22 | AMBRA1 | *Ambra1* | 9.0E-16 | 8.8 |
| B1ASB6 | SPOCD1 | *Spocd1* | 4.6E-05 | 8.2 |
| Q5SYL3 | KIAA0100 | *Kiaa0100* | 4.1E-09 | 8.2 |
| Q91XW8 | FKBP6 | *Fkbp6* | 1.6E-09 | 8.2 |
| Q6VY05 | DND1 | *Dnd1* | 1.9E-55 | 7.9 |
| Q99P31 | HSPBP1 | *Hspbp1* | 3.0E-08 | 7.8 |
| Q5FW91 | TUBA3A | *Tuba3a* | 2.0E-05 | 7.7 |
| Q3UHI3 | POLR1A | *Polr1a* | 8.5E-09 | 7.6 |
| Q8CFI7 | POLR2B | *Polr2b* | 7.7E-14 | 7.3 |
| Q3TN35 | SGTA | *Sgta* | 6.2E-26 | 7.2 |
| Q99L04 | DHRS1 | *Dhrs1* | 1.0E-03 | 7.2 |
| Q9QYJ0 | DNAJA2 | *Dnaja2* | 1.1E-26 | 7.1 |
| Q8VD46 | ASZ1 | *Asz1* | 1.7E-09 | 7.0 |
| Q3UF95 | BAG6 | *Bag6* | 1.3E-05 | 7.0 |
| Q6P9Q2 | POLR1B | *Polr1b* | 3.2E-05 | 6.6 |
| Q5NTY0 | DNAJA1 | *Dnaja1* | 9.9E-22 | 6.4 |
| P01798 | HVM29 | *Hvm29* | 1.1E-03 | 6.1 |
| B7ZNX2 | ANAPC1 | *Anapc1* | 1.2E-07 | 5.9 |
| Q792E4 | PFDN6 | *Pfdn6* | 6.5E-05 | 5.8 |
| A0A075B666 | IGKV12-47 | *Igkv12-47* | 1.6E-03 | 5.8 |
| Q7TMG8 | GBAS | *Gbas* | 1.9E-07 | 5.6 |
| Q3TEZ2 | WDR6 | *Wdr6* | 1.5E-04 | 5.5 |
| Q6PAV2 | HERC4 | *Herc4* | 6.2E-04 | 5.5 |
| E9Q7C4 | PRR14L | *Prr14l* | 1.5E-05 | 5.5 |
| D3YY91 | LRR1 | *Lrr1* | 4.3E-07 | 5.3 |
| A0A0R4J0V5 | POLR2A | *Polr2a* | 3.1E-04 | 5.1 |
| A6H663 | BAG3 | *Bag3* | 7.8E-05 | 4.8 |
| Q8C3S2 | TANGO6 | *Tango6* | 5.5E-09 | 4.7 |
| F8WJK8 | ST13 | *St13* | 5.3E-04 | 4.4 |
| Q3THK3 | GTF2F1 | *Gtf2f1* | 7.6E-06 | 4.2 |
| Q3UHZ3 | DNMT1 | *Dnmt1* | 1.1E-03 | 4.1 |
| B2RXW7 | C4B | *C4b* | 5.3E-04 | 4.0 |
| O35465 | FKBP8 | *Fkbp8* | 1.4E-18 | 3.9 |
| K7TRH5 |  |  | 1.0E-06 | 3.8 |
| Q9DC40 | TELO2 | *Telo2* | 7.0E-05 | 3.8 |
| Q91V47 | DXS254E | *Dxs254e* | 3.2E-04 | 3.7 |
| Q3U9G2 | HSPA5 | *Hspa5* | 1.7E-29 | 3.7 |
| Q6ZQ91 | IFT140 | *Ift140* | 2.0E-05 | 3.6 |
| G3UXX5 | POLR1C | *Polr1c* | 2.7E-05 | 3.5 |
| A0A075B5J9 | IGKV17-127 | *Igkv17-127* | 9.0E-04 | 3.4 |
| E0CZ27 | H3F3A | *H3f3a* | 1.6E-03 | 3.4 |
| Q99MV1 | TDRD1 | *Tdrd1* | 1.3E-17 | 3.3 |
| Q544Z7 | APEX1 | *Apex1* | 8.8E-04 | 3.3 |
| O55222 | ILK | *Ilk* | 3.2E-04 | 3.1 |
| Q0VF62 | BCAS3 | *Bcas3* | 3.2E-05 | 3.1 |
| A2NVF0 | IGKV10-96 | *Igkv10-96* | 1.7E-08 | 3.1 |
| F8WHU3 | ZYG11A | *Zyg11a* | 8.8E-04 | 3.0 |
| Q3UFV9 | DNAJC3 | *Dnajc3* | 8.6E-04 | 3.0 |
| A2AQW0 | MAP3K15 | *Map3k15* | 6.4E-05 | 3.0 |
| Q8N7N5 | DCAF8 | *Dcaf8* | 1.8E-04 | 3.0 |
| Q3U1J4 | DDB1 | *Ddb1* | 2.5E-19 | 3.0 |
| Q8CDG1 | PIWIL2 | *Piwil2* | 5.1E-14 | 2.9 |
| X5J516 | IGHV1-9 | *Ighv1-9* | 1.8E-08 | 2.9 |
| P01863 | IGHG | *Ighg* | 1.5E-09 | 2.9 |
| F6TCF9 | BAG1 | *Bag1* | 3.5E-05 | 2.8 |
| A0A075B5R8 | IGHV11-2 | *Ighv11-2* | 4.1E-04 | 2.8 |
| P01027 | C3 | *C3* | 3.4E-05 | 2.8 |
| A0A0A6YWI9 | IGHV1-11 | *Ighv1-11* | 1.0E-31 | 2.8 |
| Q811N6 | ENV | *Env* | 1.8E-08 | 2.8 |
| Q66K04 | IGH | *Igh* | 8.0E-21 | 2.7 |
| A0A075B5S1 | IGHV9-1 | *Ighv9-1* | 4.8E-09 | 2.7 |
| Q3TM92 | ANAPC4 | *Anapc4* | 5.3E-04 | 2.7 |
| A0A0A6YX19 | IGHV6-4 | *Ighv6-4* | 1.1E-03 | 2.7 |
| Q542D6 | SPTLC2 | *Sptlc2* | 2.2E-04 | 2.7 |
| Q3UMJ4 | GTF2F2 | *Gtf2f2* | 4.4E-12 | 2.6 |
| Q8BYY7 | EIF5B | *Eif5b* | 5.7E-06 | 2.6 |
| K7TDV0 |  |  | 2.0E-05 | 2.6 |
| Q9CXY0 | RALA | *Rala* | 1.2E-04 | 2.6 |
| I6L985 | IGH | *Igh* | 4.5E-14 | 2.5 |
| Q9D1H7 | GET4 | *Get4* | 1.6E-05 | 2.5 |
| A0A0G2JGT0 | IGKV8-16 | *Igkv8-16* | 6.8E-12 | 2.5 |
| P39098 | MAN1A2 | *Man1a2* | 2.3E-07 | 2.5 |
| Q91Z05 | IGHG | *Ighg* | 4.1E-32 | 2.4 |
| A0A0G2JDE1 | IGHV8-12 | *Ighv8-12* | 1.1E-05 | 2.4 |
| P05213 | TUBA1B | *Tuba1b* | 5.8E-28 | 2.4 |
| Q9JLN9 | MTOR | *Mtor* | 5.3E-04 | 2.4 |
| Q9QZD8 | SLC25A10 | *Slc25a10* | 6.1E-06 | 2.3 |
| Q3TXV1 | PSMD2 | *Psmd2* | 9.9E-08 | 2.3 |
| Q546G4 | ALB | *Alb* | 5.8E-04 | 2.2 |
| Q9D037 | ATP5L | *Atp5l* | 1.9E-10 | 2.2 |
| P63038 | HSPD1 | *Hspd1* | 1.0E-12 | 2.2 |
| X5J4F8 | IGHV14-2 | *Ighv14-2* | 2.0E-14 | 2.2 |
| Q8BN87 | MDK | *Mdk* | 4.6E-04 | 2.1 |
| K7U630 |  |  | 7.2E-16 | 2.1 |
| P99024 | TUBB5 | *Tubb5* | 2.6E-19 | 2.1 |
| A0A140T8N5 | IGKV6-23 | *Igkv6-23* | 1.7E-05 | 2.1 |
| A0A0E4B366 | IGHG2C | *Ighg2c* | 1.8E-09 | 2.1 |
| Q3UZ17 | LUC7L2 | *Luc7l2* | 6.9E-09 | 2.0 |
| P01629 |  |  | 5.7E-06 | 2.0 |
| A0A1E1GJG6 | GM5571 | *Gm5571* | 6.5E-05 | 2.0 |

**Supplementary Table 3.**
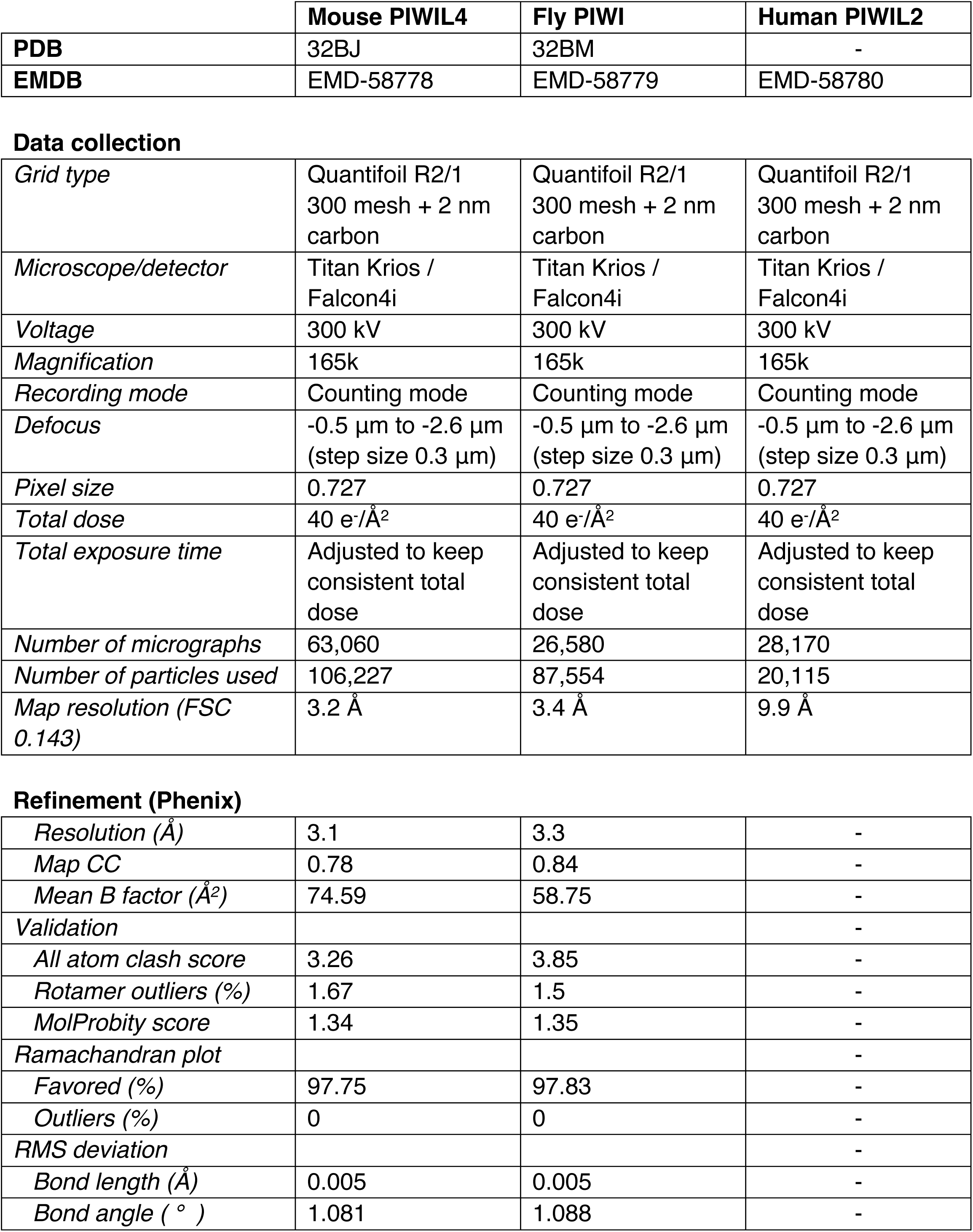
Cryo-EM data collection, refinement and validation statistics.

|  | Mouse PIWIL4 | Fly PIWI | Human PIWIL2 |
| --- | --- | --- | --- |
| **PDB** | 32BJ | 32BM | - |
| **EMDB** | EMD-58778 | EMD-58779 | EMD-58780 |
| Data collection |  |  |  |
| *Grid type* | Quantifoil R2/1<br>300 mesh + 2 nm<br>carbon | Quantifoil R2/1<br>300 mesh + 2 nm<br>carbon | Quantifoil R2/1<br>300 mesh + 2 nm<br>carbon |
| *Microscope/detector* | Titan Krios /<br>Falcon4i | Titan Krios /<br>Falcon4i | Titan Krios /<br>Falcon4i |
| *Voltage* | 300 kV | 300 kV | 300 kV |
| *Magnification* | 165k | 165k | 165k |
| *Recording mode* | Counting mode | Counting mode | Counting mode |
| *Defocus* | -0.5 $\mu\text{m}$ to -2.6 $\mu\text{m}$<br>(step size 0.3 $\mu\text{m}$) | -0.5 $\mu\text{m}$ to -2.6 $\mu\text{m}$<br>(step size 0.3 $\mu\text{m}$) | -0.5 $\mu\text{m}$ to -2.6 $\mu\text{m}$<br>(step size 0.3 $\mu\text{m}$) |
| *Pixel size* | 0.727 | 0.727 | 0.727 |
| *Total dose* | 40 $\text{e}^-/\text{\AA}^2$ | 40 $\text{e}^-/\text{\AA}^2$ | 40 $\text{e}^-/\text{\AA}^2$ |
| *Total exposure time* | Adjusted to keep<br>consistent total<br>dose | Adjusted to keep<br>consistent total<br>dose | Adjusted to keep<br>consistent total<br>dose |
| *Number of micrographs* | 63,060 | 26,580 | 28,170 |
| *Number of particles used* | 106,227 | 87,554 | 20,115 |
| *Map resolution (FSC*<br>*0.143)* | 3.2 $\text{\AA}$ | 3.4 $\text{\AA}$ | 9.9 $\text{\AA}$ |

|  |  |  |  |
| --- | --- | --- | --- |
| *Resolution ($\text{\AA}$)* | 3.1 | 3.3 | - |
| *Map CC* | 0.78 | 0.84 | - |
| *Mean B factor ($\text{\AA}^2$)* | 74.59 | 58.75 | - |
| *Validation* |  |  | - |
| *All atom clash score* | 3.26 | 3.85 | - |
| *Rotamer outliers (%)* | 1.67 | 1.5 | - |
| *MolProbity score* | 1.34 | 1.35 | - |
| *Ramachandran plot* |  |  | - |
| *Favored (%)* | 97.75 | 97.83 | - |
| *Outliers (%)* | 0 | 0 | - |
| *RMS deviation* |  |  | - |
| *Bond length ($\text{\AA}$)* | 0.005 | 0.005 | - |
| *Bond angle (°)* | 1.081 | 1.088 | - |

